# hECA: the cell-centric assembly of a cell atlas

**DOI:** 10.1101/2021.07.21.453289

**Authors:** Sijie Chen, Yanting Luo, Haoxiang Gao, Fanhong Li, Jiaqi Li, Yixin Chen, Renke You, Minsheng Hao, Haiyang Bian, Xi Xi, Wenrui Li, Weiyu Li, Mingli Ye, Qiuchen Meng, Ziheng Zou, Chen Li, Haochen Li, Yangyuan Zhang, Yanfei Cui, Lei Wei, Fufeng Chen, Xiaowo Wang, Hairong Lv, Kui Hua, Rui Jiang, Xuegong Zhang

## Abstract

Single-cell omics data can characterize multifaceted features of massive cells and bring significant insights to biomedical researches. The accumulation of single-cell data provides growing resources for constructing atlases for all cells of a human organ or the whole body. The true assembly of a cell atlas should be cell-centric rather than file-centric. We proposed a unified information framework enabling seamless cell-centric data assembly and developed a human Ensemble Cell Atlas (hECA) as an instance. hECA version 1.0 assembled scRNA-seq data across multiple studies into one orchestrated data repository. It contains 1,093,299 labeled cells and metadata from 116 published human single-cell studies, covering 38 human organs and 11 systems. We invented three methods of applications based on the cell-centric assembly: “*In data*” cell sorting enables targeted data retrieval in the full atlas with customizable logic expressions; The “quantitative portraiture” system provides a multi-view presentation of biological entities (organs, cell types, and genes) of multiple granularities; The customizable reference creation allows users to use the cell-centric assembly to generate references for their own cell type annotations. Case studies on agile construction of user-defined sub-atlases and “*in data*” investigation of CAR-T off-targets in multiple organs showed the great potential of cell-centric atlas assembly.

## INTRODUCTION

Cells are the basic structural and functional units of the human body. Different types of cells that reside in different tissues and organs of the human body could be characterized by their various molecular features, especially transcriptomic features. Building molecular atlases at single-cell resolution of all cell types in the human body in health or disease can provide basic references for future biomedical studies. The HCA (Human Cell Atlas) and the HuBMAP (Human BioMolecular Atlas Program) (Regev et al., 2017; Snyder et al., 2019) are two major efforts for building such references, among several other projects aimed at similar or related goals. These big consortiums have involved labs worldwide in generating and organizing data (Arazi et al., 2019; Azizi et al., 2018; Bayraktar et al., 2020; Chevrier et al., 2017; Cillo et al., 2020; Corridoni et al., 2020; Fernandez et al., 2019; Grubman et al., 2019; Vieira Braga et al., 2019; Villani et al., 2017; Wang et al., 2020a).

The rapid development and democratization of single-cell technologies has propelled a wave of single-cell studies. Massive amounts of single-cell transcriptomic data were pouring to the public. The data from these studies have covered all major adult human organs (e.g., Aizarani et al., 2019; Bayraktar *et al*., 2020; Guo et al., 2018; Han et al., 2020; Litviňuková et al., 2020; Pellin et al., 2019), key developmental stages (e.g., Asp et al., 2019; Cao et al., 2020; Cui et al., 2019; Guo et al., 2020; Kernfeld et al., 2018; Park et al., 2020; Zhong et al., 2018), samples from healthy donors and disease patients (e.g., Grubman *et al*., 2019; Reyfman et al., 2019; Wang *et al*., 2020a; Zhang et al., 2020). Most single-cell studies have generated data for their specific scientific questions rather than for building atlases. But these scattered public single-cell data suggests an alternative approach of building cell atlases in a bottom-up “shot-gun” manner if data can be assembled from multiple sources.

Assembling data of massive amount cells from multiple sources into an ensemble atlas, rather than just collecting and indexing the data files from the sources, has many technical and conceptual challenges (Chen et al., 2021b). Firstly, single-cell omics data describes the abundances and occurrences of a large variety of molecules and molecular events in many single cells. The data dimensionality and volume requires very wide and long sample-by-feature tables for storage. Traditional relational databases fail to hold data of such sizes. Special infrastructure adaptable for storage and efficient retrieval of single-cell data is needed. Secondly, a universal indexing scheme for cells in the human body is lacking. At the macroscopic level, cells can be indexed by their anatomic and spatial arrangements such as organs and regions. But the microscopic location of each single cell is not determined or destined. There can be multiple factors or properties that may be used to index the cells at different granularity for different study purposes. It is not feasible trying to form one fixed coordinate system to index all cells in an atlas. In addition, current annotation of cell type labels in the literature are not consistent. A standard vocabulary system for fine-grained cell identity annotation is still lacking. A unified information framework is needed to tackle these challenges (Chen *et al*., 2021b).

We developed human Ensemble Cell Atlas (hECA) as an instance of such unified information framework. In hECA v1.0, we collected the single cell transcriptomic data of 1,093,299 cells from 116 published datasets, covering 38 human organs and 146 cell types. hECA realized the cell-centric assembly of these data into a unified data repository with a special storage engine we called uGT or unified Giant Table. It has the capacity to contain all possible attributes that could be used as indexes of the cells besides the transcriptomic data. The “assembly” of a cell atlas is the unified storage and organization of all the information, rather than an ordering of the cells with a fixed coordinate system. Such cell-centric assembly allows for multiple ways of indexing the cells in the atlas. Along with uGT is a unified Hierarchical Annotation Framework (uHAF) we developed for hECA. Annotating with uHAF made cell type labels from different datasets comparable and consistent. We also developed an API named ECAUGT (pronounced “e-caught”) for efficient retrieval of cells in the atlas. With these technologies, we developed three new schemes for comprehensive use of the assembled atlas: (1) “*in data*” cell sorting for selecting cells from the virtual human body of the assembled cells using flexible combinations of logic expressions, (2) a “quantitative portraiture” system for representing the full information of genes, cell types, and organs, and (3) “customizable reference creation” for users to customize their own references for cell type annotation tasks. Case examples on the agile construction of specific sub-atlases and *in data* investigation of drug of-targets throughout the whole body showed that the hECA opens many new possibilities in biomedical research using the assembled cell atlas.

## RESULTS

### Overview of hECA v1.0

Unlike genomes, elements in cell atlases cannot be indexed or arranged in a simple linear order or in a deterministic 3D coordinate system. There are many possible ways of logical arrangements of cells at multiple granularities. The assembly of a cell atlas should convey the multifaceted nature of the data and allow users to search with customized conditions between different indexing methods.

We reasoned that the ideal cell atlas assembly should be: all cells and their multifaceted indexing coordinates should be deposited in one data system; the data system should support flexible searching using any indexing criteria thus enabling viewing and utilizing the atlas at multiple possible angles and resolutions. The system should be “cell-centric” in the sense that cells rather than datasets or files are the basic unit of data deposit, organization and retrieval.

We developed such a system called human Ensemble Cell Atlas or hECA by assembling single-cell RNA-seq data collected from scattered literature. The data origins include large projects such as the Human Cell Landscape (Han *et al*., 2020) and Allen Brain Atlas (Sunkin et al., 2013), as well as smaller datasets in many other publications (details of the data sources are given in **Table S1)**. We collected and processed the data as described in **STAR Methods**. The current version (hECA v1.0) contains data of 1,093,299 cells covering 38 human organs and 11 systems (integumentary, endocrine, urinary, cardiovascular, lymphatic, nervous, respiratory, digestive, muscular, reproductive, and skeletal systems). All cells were annotated with a unified framework of 146 cell type labels (see **Data S1**). **Table 1** summarizes the numbers of collected cells in each organ.

**Table 1.** Summary of cells collected in the organs in hECA v1.0.

| Table 1. Summary of cells collected in the organs in hECA v1.0 |  |  |  |  |  |
| --- | --- | --- | --- | --- | --- |
| # | Organ | # of cells | # | Organ | # of cells |
| 1 | Adipose | 1,362 | 20 | Oesophagus | 87,947 |
| 2 | Adrenal gland | 15,065 | 21 | Ovary | 6,927 |
| 3 | Bladder | 3,980 | 22 | Pancreas | 26,566 |
| 4 | Blood | 29,514 | 23 | Placenta | 9,926 |
| 5 | Bone marrow | 8,671 | 24 | Pleura | 19,695 |
| 6 | Brain | 214,314 | 25 | Prostate | 2,445 |
| 7 | Bronchi | 12,553 | 26 | Rectum | 5,718 |
| 8 | Colon | 22,919 | 27 | Rib | 5,907 |
| 9 | Duodenum | 3,743 | 28 | Skin | 6,618 |
| 10 | Eye | 47,275 | 29 | Spinal cord | 4,483 |
| 11 | Gallbladder | 14,733 | 30 | Spleen | 15,806 |
| 12 | Heart | 210,597 | 31 | Stomach | 22,187 |
| 13 | Ileum | 3,132 | 32 | Testis | 13,210 |
| 14 | Intestine | 41,851 | 33 | Thymus | 4,516 |
| 15 | Jejunum | 4,198 | 34 | Thyroid | 12,599 |
| 16 | Kidney | 45,368 | 35 | Ureter | 2,205 |
| 17 | Liver | 26,475 | 36 | Uterine tube | 6,496 |
| 18 | Lung | 90,521 | 37 | Uterus | 8,096 |
| 19 | Muscle | 26,029 | 38 | Vessel | 9,652 |

The overall conceptual structure of hECA is illustrated in **Figure 1**. It is an instance of the ideal unified information framework that are required for cell atlas assembly (Chen et al, 2021b). It composes of three key components: a unified Giant Table (uGT), a unified Hierarchical Annotation Framework (uHAF) and an API ECAUGT for retrieving data. uGT is a unified storage system that is technically unbounded in both rows and columns for future increases of cell numbers and feature dimensions. The basic storage of hECA is flattened to a millions by billions giant table (designed scale, in hECA v1.0 it’s 43,878 by 1,093,299). All features and metadata (any related information such as tissue origin, donor description, data source, etc. see **STAR Methods** for complete list) of every single cell are stored together. This unified storage strategy allows instant access of all information of every cell, enables flexible ways of retrieving, analyzing and comparing data, and breaks the boundary of data sources while preserving the original information. uHAF is a structured knowledge graph serving as the underlying index system for hECA. This structure organizes data into a hierarchy, provides perspectives for presenting relations and interactions between entities while preserving space for future knowledge and data growth. We provided quantitative portraits of all existing entities on this structure and a tree-view filter of the structure for cell sorting. ECAUGT is a multi-functional API (application programming interface) for manipulating data in hECA. Based on it we built hECA as a highly interactive system with both graphical user interface and command line tools. Users can access both data and structured annotations with these interfaces for downstream applications. The web interface has provided useful tools for browsing, visualizing, summarizing and analyzing pre-selected or user-selected data in hECA. Advanced users can write codes with the API for more sophisticated re-organization and deeper analyses of the data. Details of uGT, uHAF and ECAUGT are given in **STAR Methods**.

**Figure 1.**
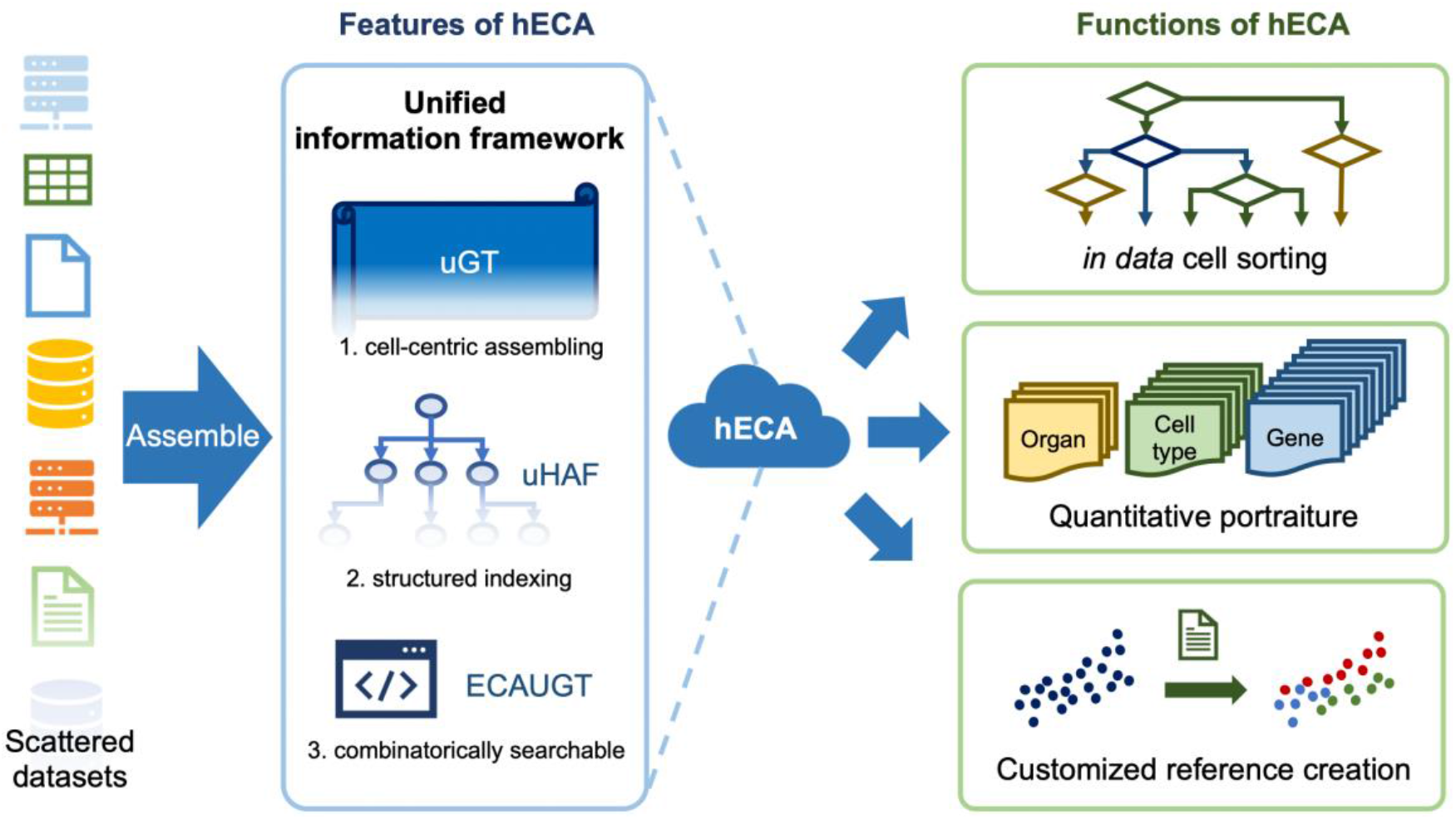
Overview of hECA. Scattered data are assembled into the ensemble cell atlas using a unified information framework. The framework includes uGT, uHAF and ECAUGT. They made hECA the first cell-centric assembled cell atlases with structured indexing and support for combinatorial searching. Based on these features, three novel functions were built on hECA: “*in data”* cell sorting, quantitative portraiture and customizable reference creation.

Based on these technologies and the assembled data, we invented three novel ways of using cell atlases for comprehensive biomedical investigations. We developed an “*in data*” cell sorting technology that takes the assembled atlas as a virtual human body to select cells from with advanced logic conditions. We developed a “quantitative portraiture” system for representing biological entities involved in the atlas from multiple angles in a holographic manner instead of only using a few marker genes. For the basic application of using cell atlas data to annotate users’ in-house data, we provided the feature of customizable reference creation. Users can define their own logic combinations to select and organize cells in hECA to form the reference for their specific queries.

### *In data* cell sorting enables comprehensive virtual cell experiments as a new research paradigm

Cell sorting is a fundamental technique in cell biology. The “*in data*” cell sorting is an innovative virtual cell experiment scheme we introduced in hECA facilitated by cell-centric organization of data. *In data* cell sorting allows users to select any cell of interest in the atlas according to any feature of the cell. When the data in the atlas provides sufficient coverage on all major tissues, organs and cell types of the human body, the cell-centric assembled cell atlas becomes a virtual human body. To precisely pinpoint the required cells from the virtual body, the criteria are expressed as a combination of logic expressions, such as desired expression range of one or multiple genes, required organs, tissue origins or developmental stages, donor’s gender and age, etc. This sorting scheme has higher flexibility, resolution and finer granularity than traditional cell sorting on *in vivo* or *in vitro* samples. The sorting dimensions is not restricted by several surface markers as for flow cytometry, but cab be extended to precisely measuring tens of thousands of features. The source materials for the sorting is not restricted by samples collected in one study, but extended to all cells with desired properties from various studies in the whole atlas. Designing cell experiments becomes a matter of writing a code of logic expression for searching hECA. This opens the new paradigm in cell biology: *in data* cell sorting followed by *in silico* computational experiments. This “*in data* experiment” paradigm will facilitate scientists to conduct investigations in the data space beyond the limitations in traditional *in vivo* or *in vitro* experiments.

*In data* cell sorting can be implemented on the hECA interactive web interface or using the Python package ECAUGT. Here we show an example of the sorting: to sort for all T cells in the heart with normalized expression of gene PTPRC greater than 0.5 and that of CD3D or CD3E greater than 0.5, users can simply type the logic expression in python:

~~~
rows_to_get = ECAUGT.query_cells(“organ==Heart && cell_type == T cell”,
include_children=TRUE)
gene_condition = ECAUGT.seq2filter(“PTPRC > 0.5 && (CD3D>=0.5 || CD3E>=0.5)”)
ECAUGT.get_columnsbycell_para(rows_to_get = rows_to_get, cols_to_get=[‘CD3E’,’PTPRC’],
col_filter=gene_condition
~~~

and hECA will return the selection results (of around 210,000 cells in current version) in about 190 seconds.

This example shows the logical clarity, convenience and efficiency of *in data* cell sorting. By contrast, the typical cell sorting workflow composed of multiple filtering steps is more complicated. To obtain the regulatory T cells (Treg) from certain type of human tissue sample, a researcher needs to use the marker protein PTPRC (also known as CD45) to distinguish immune cells (PTPRC+) from other lineages of cells (PTPRC-), use CD3 to select the T cells (PTPRC+ and CD3+) from the PTPRC+ cells, and then use CD4, IL2RA (also known as CD25), and FoxP3 markers to filter out other T cells and get the Treg cells. The types of cells that can be selected depend on the availability and identifiability of surface markers of the cells under study, and the discriminating power of the flow cytometry technology. This sorting practice is much lengthy and time-consuming than the *in data* sorting. And *in data* sorting can apply many selection criteria that may not be possible for flow cytometry. With growing coverage of hECA, researchers can conduct all kinds of pre-experiments with *in data* cell sorting to accelerate research loop.

Another advantage of *in data* cell sorting is swift multi-step iteration. Users can jump back and forth in sorting steps to make comparison and achieve optimal result. They can adjust sorting criteria based on analysis of previous steps, without bother waiting for another experiment loop. For users to have a quick overview of sorted cells, we provided a real-time analysis function on the web interface. The real-time analysis includes the following properties of the selected cell group: 1) cell type composition in all and every organ; 2) expression distribution of interested gene across cell types and organs; 3) pseudo-FACS simulating a real FACS to show relative expression level between any two interested genes. Users can conduct next step of cell sorting based on the results of real-time analysis, without the trouble of downloading and locally analyzing the whole dataset. We provided five examples of utilizing *in data* cell sorting, three of them are done with web interface, the other two are shown in ECAUGT with vignette and detailed explanations (see **STAR Methods**).

We conducted two case examples on leveraging the potential of *in data* cell sorting: 1) agile construction of atlases of particular cell types; 2) off-target prediction of targeted therapy. These cases demonstrated in details of how hECA can be used to conduct comprehensive studies of cells across the human body in an unprecedented way.

### Case study 1: agile construction of a draft T-cell metabolic landscape

In the first case example, we built a draft T-cell sub-atlas to show the power of hECA in agile construction of cell landscapes across studies. This case also shows how to compare the metabolic activity heterogeneities between different organs/cell types in a high-throughput way from the public data.

T lymphocyte is an essential cell type in human immune system. They adapt to multifarious microenvironments as they circulate through or reside in human body. Their differentiation, activation and quiescence are regulated by diverse metabolites in local microenvironment (Buck et al., 2015; Chapman et al., 2020; Shyer et al., 2020; Yin et al., 2019). Recent studies reported that microbial bile acid metabolites promoted the generation of regulatory T cells in the intestine, which is associated with inflammatory bowel disease (IBD) (Campbell et al., 2020; Hang et al., 2019; Song et al., 2020), suggesting that targeting metabolic pathways of T cell activation and differentiation may improve therapeutic outcomes of IBD patients (Li et al., 2021). Comprehensive survey of the metabolism of T cells across multiple organs is crucial for better understanding intrinsic responses of T cells to microenvironment changes, but *in vivo* or *in vitro* experiments on multiple organs are not easy. Xiao et al proposed a computational pipeline to study the metabolic landscape of cells from single-cell transcriptomic data (Xiao et al., 2019). The cell-centric assembly of cells of all types in all organs in hECA allowed us to conduct *in data* study on T cell metabolism across all organs, instead of searching through datasets scattered in the literature.

Using ECAUGT, we first sorted all cells in uGT with label “T cell” and associated names (such as “CD4 T cell”, “CD8 T cell”, “Activated T cell”, etc.) across all organs (**Figure S1A**). To include cells that might be annotated to other cell types, we also searched for cells with normalized expression values of PTPRC, CD3D or CD3E greater than 0.5 across all organs (**Figure S1B, S1C**). Then we filtered the cells by the expression of a list of negative markers such as COL1A1, CD79A (the full list provided in **Table S4**). We conducted clustering analysis on cells from the same organs, and obtained a series of candidate clusters in each organ (**Figure 2A**). We removed clusters with low expression levels of CD3D, CD3E or CD3G as they are unlikely to be T cells. After these steps in hECA, we built an agile cell atlas of T cells across 18 organs (lung, pancreas, blood, liver, muscle, thymus, jejunum, rectum, colon, kidney, gallbladder, stomach, thyroid, intestine, spleen, bone marrow, eye, and vessel).

**Figure 2.**
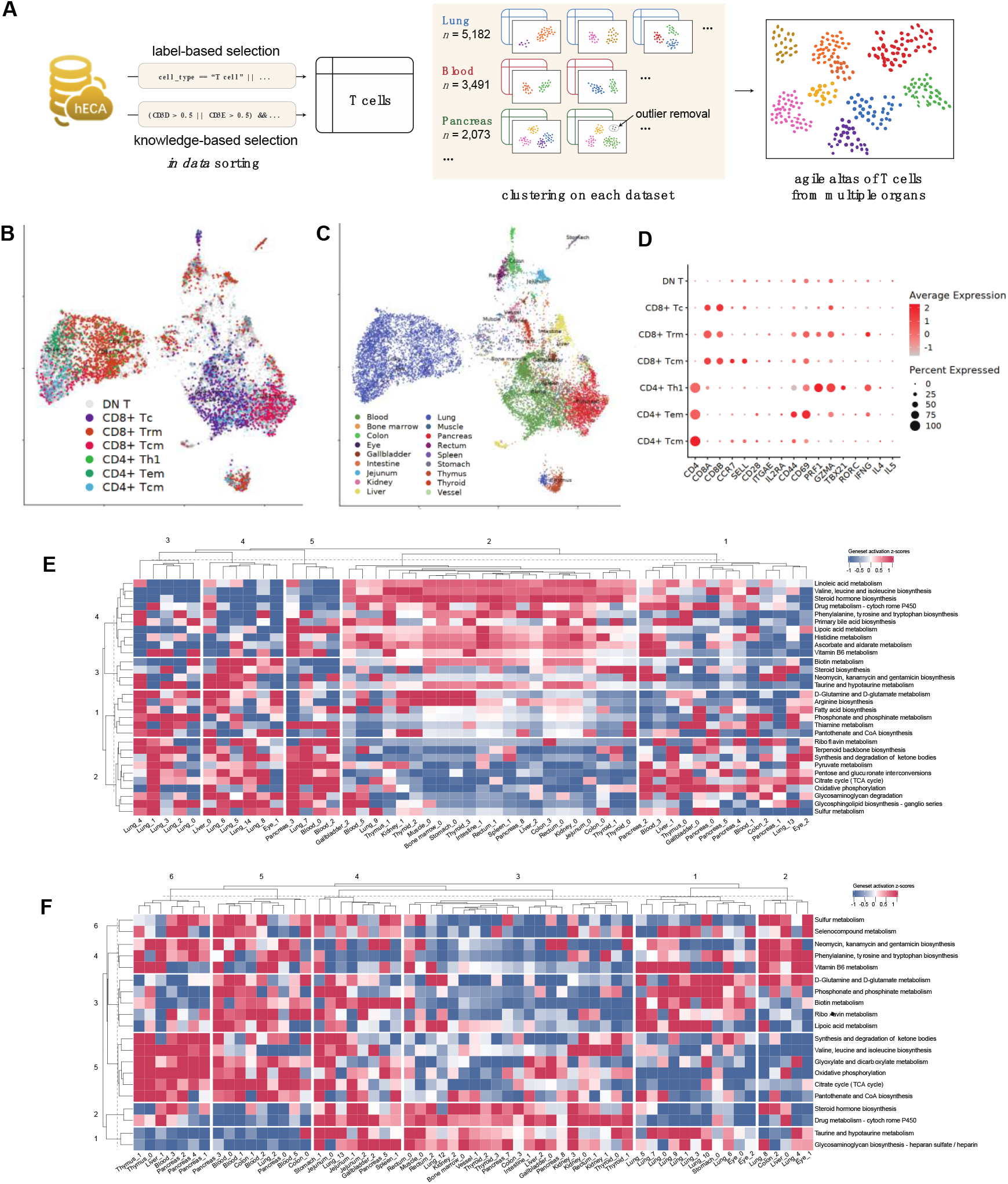
The agile creation of a draft T-cell metabolic landscape across multiple organs from hECA. (A) Workflow of the *in data* cell sorting from hECA to build the agile T cell atlas. (B) Subtypes of selected T cells displayed on UMAP. DN T: Double negative T cell, CD8+ Tc: CD8+ Cytotoxic T cell, CD8+ Trm: CD8+ resident memory T cell, CD4+ Th1: CD4+ T helper cell type 1, CD4+ Tem: CD4+ effector memory T cell, CD4+ Tcm: CD4+ central memory T cell. (C) Organ origins of selected T cells organ origin displayed on UMAP. (D) Gene expression signatures of the identified T cell subtypes. (E-F) Heatmaps showing z-scores of activity scores of major metabolic pathways of the T cell subtypes in multiple organs. (E) for CD4+ T cells and (F) for CD8+ T cells. Each row in the heatmap corresponds to one selected term in the KEGG metabolism pathway database, and each column corresponds to one T cell subcluster.

The following experiment are downstream analysis performed out of hECA to prove the viability of the constructed T cell atlas. To assign accurate annotations to the cells in the T cell atlas, we performed hierarchical clustering using signature genes CD4, CD8A and CD8B, and divided the cells into 6 subgroups of 3 major groups (**Figure S2A**). The three major groups are CD4+, CD8+ and double-negative (CD4- and CD8-) T cells (**Figure S2B**). For the CD4+ and CD8+ groups, we further annotated the cells as resident memory T cells, central memory T cells, effector memory T cells, naïve T cells, cytotoxic T cells, etc. according to the positive markers listed in **Table S5. Figures 2B, 2C** show the UMAP of the CD4+ and CD8+ T cells with the subtype annotations and with the organ origin of the cells, respectively. **Figure 2D** shows the gene expression signatures of the identified T cell subtypes. For the double-negative cluster, we marked them as “T cells” without further analysis as there might be cells false negatives in CD4 or CD8 expression due to possible dropout events in scRNA-seq data.

For a sketchy study on the metabolic landscape of T cells across multiple organs, we evaluated each cell’s metabolic activity scores with GSVA, which produced comparable values across multiple clusters or datasets and alleviated possible batch effects in the data from multiple sources (Hänzelmann et al., 2013). The genes of the metabolic pathways are derived from KEGG (Kanehisa et al., 2020) and Xiao et al.’s work (Xiao *et al*., 2019). A heatmap of the obtained draft metabolic landscape of T cells of their activity scores of all major metabolic pathways across the human body are shown in **Figures 2E** and **2F**. Such landscapes can help to reveal different metabolic patterns across organs. For example, we found organ-level metabolic variations in lungs from the metabolic activities of organ-level CD4+ T cell clusters in **Figure 2E** and those of the organ-level CD8+ T cell clusters in **Figure 2F**. For CD4+ T cells, we observed lung-enriched metabolic pathway activations in the pathways of riboflavin metabolism, terpenoid backbone biosynthesis, TCA cycle, oxidative phosphorylation, sulfur metabolism, and D-Glutamine and D-glutamate metabolism (row blocks 1 & 2 of the lung-origin T cell clusters in **Figure 2E**). Similar enrichments can also be observed in the lung-origin CD8+ T cell clusters in **Figure 2F**.

These observations deserve further investigations. They showcased the potential of cross-organ in data cell experiments enabled by hECA which are otherwise hard to conduct in traditional experiment settings.

### Case study 2: *in data* discovery of side effects in targeted therapy

In the second case example, we utilized *in data* cell sorting to investigate possible off-target effects in cancer therapy. This case study shows hECA’s potential application in investigating human diseases.

A great part (∼97%) of cancer drugs tested in clinical trials failed to get approval from FDA, mainly due to their insufficient efficacy or unexpected toxicities to organs where drugs were not designed to take effect (Lin et al., 2019). Off-target effects are usually not easy to observed on animal models. Prediction of cellular toxicities across the whole body can significantly reduce improper clinical trials and increase efficiency of new drugs discoveries. This is a typical scenario where we should conduct *in data* experiment on the virtual human body of cells to test drugs before clinical trials on human patients.

In previous research, computational investigation of off-target effects or neurotoxicity effects of targeted therapy took multiple steps. Researchers first chose a group of organs as suspects of side effects based on existing knowledge. They need to review the literatures to search for single-cell datasets in which cells in the suspected organs have highly-expressed target genes of the candidate drug. Then they will evaluate the effect of the drug on the phenotype of these cells and therefore on the phenotype of the organs. This is a typical setting of traditional “meta-analysis”. Parker et al found that CD19+ mural cells in the human brain were potential off-tumor targets of CAR-T therapy in this way (Parker et al., 2020). They first noticed from previous literature that CD19 CAR-T therapy could introduce neurologic adverse reactions. Then they collected 3 single-cell datasets of human brain: prefrontal cortex (Zhong *et al*., 2018), forebrain (La Manno et al., 2018) and ventral forebrain (La Manno et al., 2016). After applying standard reprocessing on each dataset, they manually annotated cells by comparing highly enriched genes to known cell-type markers. They observed on the UMAP a small population of cells in the first dataset expressed both CD19 and CD248 (a marker for mural cells). They further identified that these cells were pericytes and verified them in all three datasets. This type of meta-analysis depends much on the existing hints or guesses on possible off-target organs and involves heavy efforts on data collection and reprocessing.

We followed the example of Parker’s work (Parker *et al*., 2020) to study the possible off-target effects of CAR-T therapy in a more automatic way using hECA. CD19 is a common target of CAR-T therapy treating B-cell lymphoma (Wei et al., 2019). Neurological toxicity is one of the major side-effects (Rubin et al., 2019). To study why this toxicity occurs and whether other organs might be affected by CAR-T therapy, we used a filtering criterion on CD19 expression for *in data* cell sorting in hECA. Totally 2,566 CD19+ cells passed the filter (**Figure 3B**). This therapy aims to target malignant B cells for curing lymphoma. But B cells and plasma B cells only compose ∼53% of the selected CD19+ cells (**Figure 3C, Figure S4, Table S6**). The other cells in the selected group include endothelial cells, microglia and neurons in the brain, cardiomyocytes and fibroblasts in the heart and lung, enterocytes in the rectum, etc. (**Figure 3D, Figure S4, Table S6**). They all have the potential of suffering from off-targets of the therapy. This result explained why encephalopathy was often observed and cells constructing vessels were targeted by the drug (Parker *et al*., 2020). Our results also suggest that there is possible toxicity on the circulatory system and digestive system, which can also be validated by reports in the literature (Yáñez et al., 2019).

**Figure 3.**
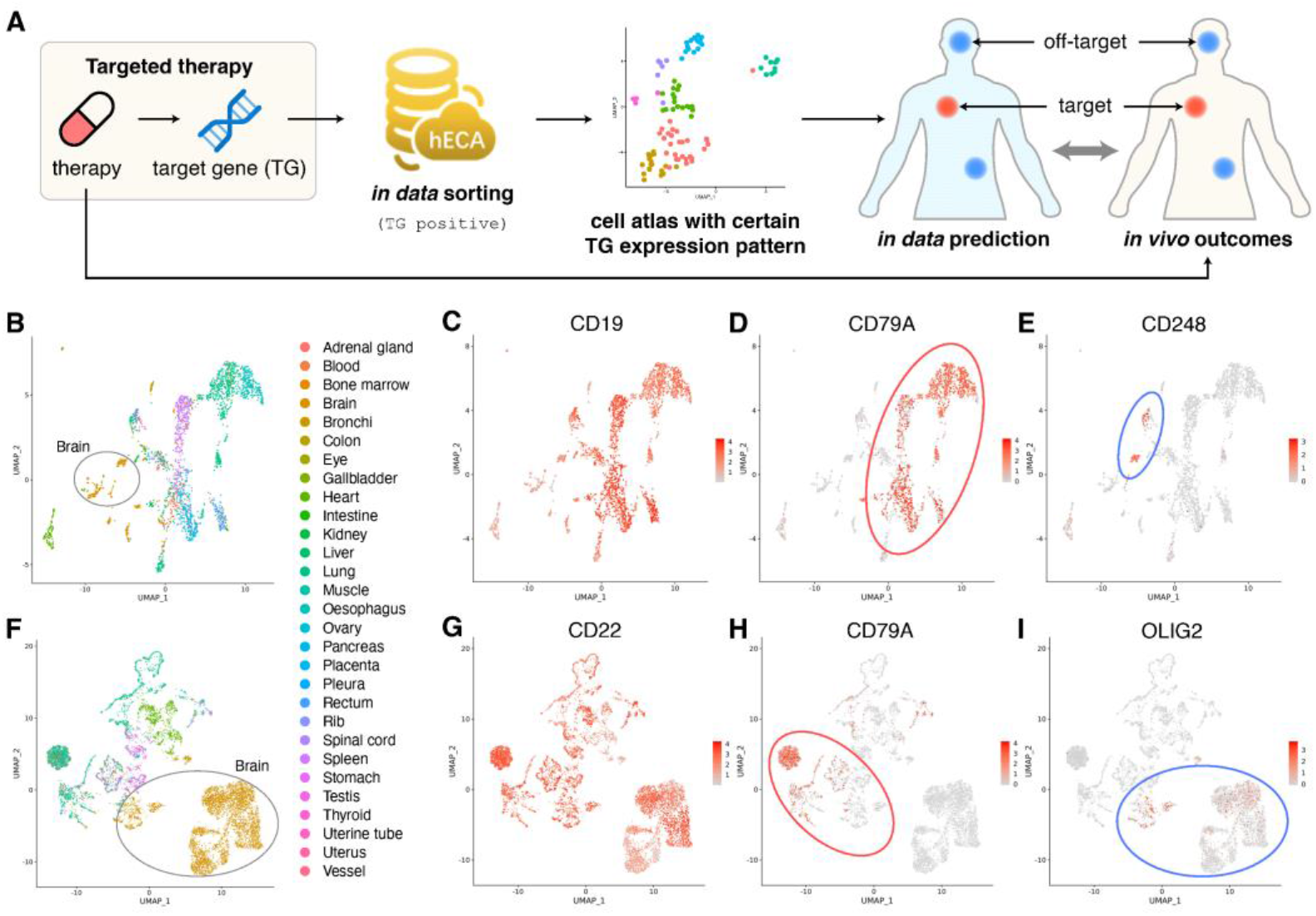
*In data* experiments with hECA facilitating discoveries of side effects of targeted drugs. (A) The diagram of using *in data* cell sorting to predict targets and off-targets of targeted therapy. Red dots and blue dots in the human body represent the target effected sites and side effected sites, respectively. The red and blue dots in the UMAP represent the treatment effected cells and side effected cells, respectively. (B) Visualization of CD19+ cells (expression>0.1) in UMAP, colored by organ origins of cells. CD19 is the target gene of the targeted therapy. (C) Visualization of CD19 expression levels of those CD19+ cells. (D) Visualization of CD79A expression levels of those CD19+ cells. CD79A is a marker for B cells. (E) Visualization of CD248 expression levels of those CD19+ cells. CD248 is a marker for pericytes. (F) Visualization of CD22+ (expression>0.1) cells in UMAP, colored by organ origin of cells. CD22 is the target gene of the targeted therapy. (G) Visualization of CD22 expression levels of those CD19+ cells. (H) Visualization of CD79A expression levels of those CD22+ cells. CD79A is a marker for B cells. (I) Visualization of OLIG2 expression levels of those CD22+ cells. OLIG2 is a marker for oligodendrocytes. The color bars in (C-E) represent expression levels of CD19, CD79A and CD248, and the color bars in (G-I) represent expression levels of CD22, CD79A and OLIG2, respectively, with colors grey to red indicating expression low to high. The red and blue ellipses in (D-E) and (H-I) line out the target affected cells and off-target affected cells, respectively.

CD22 is another popular target when designing CAR-T therapy for lymphoma (Wei *et al*., 2019). Similarly, we used *in data* cell sorting in hECA and obtained 8,724 cells with CD22 expressed (**Figure 3E**). In addition to B cells (**Figure 3F, Figure S5, Table S7**), this group contains oligodendrocytes and excitatory neurons in the brain, cardiomyocytes and fibroblasts in the heard, macrophage, mast cells and monocytes in the lung, and neutrophils in the testis, etc. (**Figure 3G, Figure S5, Table S7**). These observations provide significant clues for systematic investigation on the potential side effects of targeted therapy.

This case study provided more than an example, but a systematic approach of conducting meta-analysis with *in data* cell sorting in a more efficient and effective way based the cell-centric assembly of massive single-cell data in hECA. For any specific target gene, cells that highly express the gene can be found through *in data* cell sorting, no matter which original datasets the cells are from. A profile of cellular distribution of all major human organs that contain the found cells can be built, which highlights suspected organs that might be the off targets of the drug. Detailed analyses can be further applied on the possible effects of the drug on the phenotypes of the cells by checking on the consequences of the expression change of the target gene on downstream gene expression, signaling pathways, metabolisms, interactions with other cells, etc. Quantitative analysis then can be applied on the cell compositions and cell-cell interactions in the suspected organs to evaluate the possible physiology or pathology effects. Users can adopt this approach and apply to any target cell types they want to investigate.

### Quantitative portraiture of genes, cell types and organs

The above sections illustrated how users can explore and exploit hECA with the flexible and cell-centric *in data* cell sorting engine. To better describe whole vivid pictures of the biological entities in hECA, we further developed a “quantitative portraiture” system. The system contains a set of quantitative portraits of the biological entities, including organs, cell types, and genes for all quantifiable characteristics at multiple angles. We portrayed them in the web interface at all possible levels and aspects so that users can get a comprehensive understanding of the whole system, all elements in it and their relationships. This is an upgraded approach from the current approach of using “snapshots” of marker genes to describe a cell type. In the current version, we portrayed 38 organs, 146 cell types and 43,878 genes in hECA v1.0 with the currently available data. With the growing coverage and quality of data assembled into hECA in the future, the portraiture framework will lead to “holographic” macroscopic and microscopic views of genes, cells, tissues and organs of the human body.

In gene portraits, we showed the expression distribution of a gene in each selected organ or cell type, providing a quick overview and organ-wise or cell type-wise comparison of genes of interests. We also included basic information about the gene, links to GeneCard, NCBI, Ensemble and Wikigene pages of the gene. The design of gene portraits borrowed the idea from the “gene skyline” of ImmGen (http://rstats.immgen.org/Skyline/skyline.html), a famous project that collect immunological data and profile gene expression signatures. In the portrait page of gene *PTPRC* (**Figure S5**), for example, the basic information of the gene is firstly shown, including the gene’s full name *Protein Tyrosine Phosphatase Receptor Type C*, some of the aliases, its location on the chromosome, etc. A panel “*Known as markers of”* provides information about cell types in which the gene is highly expressed. Users can browse the distribution of the gene’s expression level, grouped by the uHAF organ tree or cell type tree. The gene portraits in hECA present several major improvements compared with the gene skyline. Firstly, a distribution instead of only mean value of expression levels is provided for each gene in each cell type or organ. Besides the function of exhibiting relative expression strength between cell groups, expression distributions show more information like the percentage of cells that express the gene, or heterogeneity within a cell type which may indicate potential sub-types. Secondly, hECA gene portraits cover a wider breadth of cell types, while the data of gene skyline are restricted in immune cells. Furthermore, hECA portraits are based on the uHAF annotation. This allows the portraits to be updated timely with the expansion of uHAF when more data are assembled.

hECA cell type portraits include the organ origin of a certain cell type, marker genes in the cell type, view of the cell type in embedding space and the position of the cell type in the uHAF tree (**Figure S6**). A cell type is mainly characterized by two types of information: organs that contain the cell type, and the expression patterns of genes that are specific to the cell type. hECA v1.0 portrayed 146 of the 416 cell types organized by the hECA hierarchy with the current data availability. On the hECA website, users can type in the name to search for a cell type or to click along the tree of cell types to display the cell type portrait. It includes the distribution of the cell type across organs, shown as the number of cells of this type collected in the organs, the list of marker genes with their characteristic expression ranges in the cell type, and a 2D PCA, UMAP or DensMAP visualization (McInnes and Healy, 2018; Narayan et al., 2021; Pearson, 1901) of the cells colored by the organ of the cells or the expression of a certain gene in the cells. hECA organ portraits include a organ’s cell type composition, embedding view of cell types in the organ, and a tree view of its position in uHAF (**Figure S7**). An organ is usually characterized by its anatomic and physiological features, but the full portraiture of an organ should include its full cellular and molecular features at multiple resolutions. The basic cellular information is the relative composition of cell types in the organ and in its different anatomical parts. The basic molecular information is the gene expression patterns in the organ as a whole and in its different parts, spatial locations, and at different physiological statuses. In the embedding viewer, we show the feature map of each gene in 2D visualization, showing the relationship of certain genes, cell types and organs. The current coverage and quality of the data are still far from fully characterizing the entities in an unbiased manner. Therefore, current portraits can only reflect information in the collected data rather than the complete biological picture. But the portraiture framework provides a comprehensive approach leading to the complete picture when more and more data are assembled into hECA.

It should be noted that most current single-cell sequencing technologies undergo some kind of cell selection before sequencing. For cells that are selected, the sampling efficiencies for different cell types are also not uniform (Baran-Gale et al., 2017; Phipson et al., 2017; Tung et al., 2017). There are many technical reasons that may cause biases in the measured gene expression values even in the same experiment, let alone across different experiments (Chen and Zheng, 2018; Miao et al., 2018; Miao and Zhang, 2016; Soneson and Robinson, 2018). Therefore, it is unrealistic to expect full portraits of genes, cell types or organs with high reliability and fidelity based on the currently available data. The hECA quantitative portraiture system provides a framework presenting the full information of biological entities, and sets a goal for future ideal cell atlases.

### Customized reference creation for automatic cell type classification

Every cell in hECA has standard identity labels chosen from the uHAF. Users can transfer these identity labels to their own datasets with well-designed classifiers. Many computational tools for automated cell type identification have emerged, as described in (Pasquini et al., 2021). These classification methods rely on a good selection of reference datasets to perform good label transfer because the cell composition in the reference data may lead to differences in annotation results. For instance, when studying hematopoietic development, feeding a classifier with a reference containing only hemocytes will reduce misclassification.

In hECA, the *in data* cell sorting web and programming interfaces can help users create customized references using flexible creation criteria. We also provided a list of pre-created reference datasets organized by organs, which is available at http://eca.xglab.tech/#/cellTypeList. Note that the current curated references from hECA may not be complete in the cell type composition due to the insufficient data coverage and biased sampling strategy.

## DISCUSSION

We presented hECA, a cell-centric assembled human cell atlas based on collection of data scattered in the literatures. hECA was empowered by a unified information framework providing structured indexes and combinatorial searching facility. The cell-centric assembly provides three novel applications for the atlas that could be difficult for file-centric data collections: 1) a new experiment paradigm “*in data”* cell sorting that enables efficient selection of cells that meet combinations of multiple logic conditions, 2) a “quantitative portraiture” system for holographic characterization of biological entities, and 3) a customizable reference generation function for automatic annotation of users’ query cells. Although the current data is far from providing sufficient and uniform coverage to major human organs, example applications based on hECA v1.0 already demonstrated the revolution that such cell-centric assembled cell atlas can bring to biomedical research beyond the possibility of single-cell studies or file-centric atlas collections.

There have been several efforts for gathering, collecting and archiving single-cell data. Those “data integrations” are at the dataset level rather than cell level: Data of cells from different studies and sub-studies are archived as separated files rather than merged into a single database; Databases are used to manage or index the metadata of the datasets instead of the individual cells. The typical way to use these resources is to find specific datasets from the list and download the corresponding data files to users’ local computer for in-house analyses. They provided useful resources for many studies. But it is not convenient or efficient if users need to utilize data across multiple datasets in a more comprehensive manner. Tasks such as evaluating the expression of a certain gene among multiple organs or studying cellular emigrant route need researchers to process dozens of datasets separately. These tasks need cell-centric assembly of data across studies and datasets. There has been no such reported effort yet for assembling massive single-cell data of multiple studies into a unified repository. The question of possible underlying information structures to organize and annotate all cells in an atlas has not been sufficiently studied. The unified information framework we developed in hECA provides promising solution for cell-centric assembly of cell atlas with existing data.

Although the number of cells in hECA v1.0 is still very small and the coverage of organs and cell types is very limited, case studies using this primary version already showed the advantage of cell-centric atlas assembly especially the power of *in data* experiments enabled by the assembly. The customizable annotation reference shows the other way of utilizing *in data* cell sorting. The proposed gene, cell type and organ portraitures provide a powerful framework for characterizing the full information of biological entities in a quantitative manner. Up to now, all single-cell data that have been ever generated for human cells are still only a tiny fraction of all human cells, and the data are also under the influence of multiple types of noises and biases. Therefore, the current portraits can only reveal properties of the collected data but cannot be expected of high fidelity for the underlying biology. However, keeping this reality in mind, users can already use these portraits as handy tools for exploring properties of genes, cell types and organs from a more complete view than traditional views. With the rapid advancement in data depth, coverage and quality, the portraits will provide multiple-scale holographic views of all biological entities in the human body.

Several upstream processing issues that are crucial for the construction of cell atlases, such as normalization and correction for possible batch effects. Non-uniform sampling of cells and of expressed genes is another issue that may poison global analyses of atlas data. In building hECA, we followed the currently widely accepted protocols for the upstream processing of collected data. We are fully aware that these issues are far from perfectly solved yet, either in our own work or in the community. But we choose to not stuck by those issues but to work on the key questions in assembling and utilizing the data. These two types of questions are orthogonal and we should not wait till the ideal solution of the low-level processing problems to study the assembly and advanced application problems. On the other hand, the assembling and utilization of hECA can help to pinpoint what downstream analyses are more sensitive to pre-processing and what are not. From the case examples, we can see that although expressions of genes measured in separated experiments are not precisely comparable due to possible batch effects, *in data* cell sorting on the rough expression of some genes can already reveal important organ-specific patterns and can help to discover organs that are more prone to side effects of targeted therapy. If necessary, more advanced reprocessing methods can be applied on the selected data and optimized for the specific downstream scientific investigation. This is more feasible than trying to find general optimal solutions in data preprocessing without a specific aim in the downstream study.

## ACKNOWLEDGEMENTS

This work is supported in part by NSFC grants 62050178, 61721003, 32000453 and 42050101. Many students have participated in the early attempts of literature data collection and early explorations of demo systems, including Tianxing Ma, Yi Liu, Yongzhen Yu, Yuehua Zhu, Zihao Jiang, Wuhan Qiu, Xinyi Ning, Qianshi Mei, Xiaosen Wei, Fangyuan Chen, Siting Li, Bin Zhang, Min Wu, Jiani Wang, Dongfang Mao and Xiaoxiao Nong. Dr. Aziz Khan helped to test some functions of the website. The work is also support by the Tsinghua-Fuzhou Institute of Data Technologies (TFIDT2021003).

## AUTHOR CONTRIBUTIONS

XZ conceptualized and designed the project. XZ, SC, HG, YL and FL designed the study. SC led the design and implementation of uGT. YL and HG designed uHAF with inputs from SC, FL and JL. YL designed the methods for data collection and led the efforts of data collection and annotation. FL and SC coordinated the construction of the data system and the collaboration of all teams. JL developed the portraiture system and all portraits based on current data. HG, FL, JL, SC and YL designed the web visualization. YC, SC, HB and MH developed the ECAUGT package. SC and MH conducted case study 1. HG, SC and MH conducted case study 2. RY, Weiyu Li, MY and FC developed the web user interface system and the interface with the uGT, under the supervision of H. Lv. YC, XX, HB, MH, Wenrui Li, CL, YC, H. Li and YZ curated the data under the supervision of YL. QM and ZZ participated in developing methods for data annotation, visualization and gene symbol unification method. KH, H. Lv and RJ participated in the conceptualization of the project and discussions on strategies of implementation. RJ, KH, LW and WW participated in many technical and strategical aspects of the study. XZ, SC, YL, FL, HG and JL wrote the manuscript, with inputs from all authors. All authors read and approved the manuscript.

## STAR*METHODS

### KEY RESOURCES TABLE

| RESOURCE | SOURCE | IDENTIFIER |
| --- | --- | --- |
| Deposited Data |  |  |
| hECA database | This paper | http://eca.xglab.tech/ |
| ECAUGT | This paper | https://pypi.org/project/ECAUGT/ |
| Software and Algorithms |  |  |
| Seurat | (Stuart et al., 2019) | https://github.com/satijalab/seurat |
| biomaRt | (Smedley et al., 2009) | https://bioconductor.org/packages/release/bioc/html/biomaRt.html |
| plotly | - | https://github.com/plotly/plotly.py |
| KEGG | (Kanehisa et al., 2020) | https://www.genome.jp/kegg/ |
| ontologyIndex | (Greene et al., 2017) | https://cran.r-project.org/web/packages/ontologyIndex/index.html |
| GSVA | (Hänzelmann et al., 2013) | https://www.bioconductor.org/packages/release/bioc/html/GSVA.html |
| Other |  |  |
| uHAF macroscope ontology | This paper | https://github.com/XuegongLab/hECA |
| uHAF microscope ontology | This paper | https://github.com/XuegongLab/hECA |
| uHAF macro-micro map | This paper | https://github.com/XuegongLab/hECA |
| uHAF marker reference | This paper | https://github.com/XuegongLab/hECA |
| HGNC gene symbol list | This paper | https://github.com/XuegongLab/hECA |
| GeneSymbolUniform toolkit | This paper | https://github.com/XuegongLab/hECA |
| CellMarker | (Zhang et al., 2018) | http://biocc.hrbmu.edu.cn/CellMarker/ |
| The Human Protein Atlas | (Uhlén et al., 2015) | https://www.proteinatlas.org/ |
| PanglaoDB | (Franzén et al., 2019) | https://panglaodb.se |

## RESOURCE AVAILABILITY

### Lead contact

Further information and requests for resources should be directed to and will be fulfilled by the Lead Contact, Xuegong Zhang.

### Materials availability

This study did not generate new unique reagents.

### Data and code availability

hECA database can be found in http://eca.xglab.tech/.

ECAUGT for database query can be found at https://pypi.org/project/ECAUGT/

## EXPERIMENTAL MODEL AND SUBJECT DETAILS

References to original studies that generated the single-cell transcriptome datasets analyzed in this work can be found in **Table S1**.

## METHOD DETAILS

### Dataset collection

In the first version of hECA (v1.0), we present an atlas of 1,093,299 cells from 116 datasets belonging to 21 published studies (Asp *et al*., 2019; Chen et al., 2021a; Cui *et al*., 2019; Gaublomme et al., 2019; Han *et al*., 2020; Kinchen et al., 2018; Lake et al., 2018; Lukowski et al., 2019; Madissoon et al., 2019; Menon et al., 2019; Parikh et al., 2019; Plasschaert et al., 2018; Renthal et al., 2018; Sunkin *et al*., 2013; Venteicher et al., 2017; Vieira Braga *et al*., 2019; Voigt et al., 2019; Wang *et al*., 2020a; Wang et al., 2020b; Zhong et al., 2020; Zhong *et al*., 2018), details provided in **Table S1**. We introduced hECA as an instance for cell-centric assembly of cell atlas by collecting all accessible human single cell data into a unified atlas, regardless of the technology, platform, researcher, study design or other factors in data generation. Toward this goal, we selected 20 peer-reviewed studies in hECA v1.0, preferably studies with high throughput in cell numbers and coverage of multiple healthy organs. These studies covered 38 organs and spanned the developmental stages from fetal to adult. In hECA v1.0 we only include transcriptomic data of healthy doners, but future versions will cover multi-omics data as well as data of disease samples.

In each of the studies, we collected the expression matrix of every dataset in the study. In addition, we collected all the descriptive information in study level and dataset level, and analysis results of the cells in the original papers. They are referred as metadata in hECA. Metadata includes the following information if available: sample organ, sample tissue, anatomical region, subregion, donor ID, donor gender, donor age or developmental stage, sequencing technology and original annotations that are the assigned cell type label of each cell in the original study. The completeness of metadata and annotations vary among datasets according to original studies.

### Processing of data matrixes

The collected datasets were processed for integration into uGT. The 116 datasets we collected in hECA v1.0 are all single-cell gene expression profiles, and every profile was transformed into gene by cell matrix, with each row representing a gene and each column representing a cell. For those expression value in log scale, we performed the value transformation back to raw values.

For the purpose of integrating data into uGT, we unified the gene names for all datasets. For datasets identifying gene with Ensembl ID, we used the R package biomaRt (Smedley *et al*., 2009) to convert Ensembl ID into gene symbol. Then the gene symbols of different datasets were unified with an in-house built toolkit: we compared gene symbols in the datasets to the list of 43,878 HUGO Gene Nomenclature Committee (HGNC) approved symbols (see “HGNC gene symbol list” in https://github.com/XuegongLab/hECA), all previous, withdrawn and alias symbols were converted into HGNC approved symbols. Genes that are in the list but not sequenced in any dataset were filled with zeros. After processing, every expression matrix was with 43,878 genes as rows.

For datasets that have cell type annotations in the original study, the original annotations were kept and stored in column “original_name” in the uGT. Regardless of the original annotations, we performed clustering analysis and annotation in each dataset with Seurat v3.2 (Stuart *et al*., 2019). We implemented a standard processing procedure for each dataset: We created a Seurat object from the expression matrix, conducted quality control to filter out genes and cells, selected variable genes, conducted normalization, scaling, dimensional reduction and cell clustering. The parameters for quality control and cell filtering were determined specifically for each dataset following the original studies or following the tutorial of Seurat. The parameter for cell clustering is determined based on the consistency with original cell clustering results. Then the analysis pipeline of Seurat was performed to get cell cluster-specific expressed genes. After the quality control, we got a total of 1,093,299 cells from 116 datasets.

### uGT: a unified giant table for assembling cell atlases

To support online “cell-centric” data assembly, we implemented the unified giant data table (uGT) using the NoSQL database technology (Stonebraker, 2010; Wang et al., 2017) to store data from multiple studies into one cloud repository. In this version of uGT implementation, we used the Tablestore distributed data storage service provided by Alibaba Cloud. The unified giant table supports storing and searching cells with dataset-associated attributes like organ, gender, donor age, study DOI number, and cell-specific features like cell type and ∼104 gene expression levels.

The key difference between uGT’s NoSQL database and the traditional databases is that uGT used column-based storage. Popular implementations of traditional SQL databases have a rigid width limit for each data item. For example, the limit on the number of columns is 1000 for Oracle™ and 4096 for MySQL™, which has already reached the theoretical upper limit (MySQL, 2021; Oracle, 2021). Obviously, the number of features of each cell exceeds this limit by several magnitudes. In addition, searching high-dimensional data is difficult because even if one or two columns are used for data selection, all columns are retrieved by the computer. However, in column-based NoSQL databases, the column retrieving activity is restricted to the associated columns, which greatly promotes the searching efficiency, although the insertion and update of data become difficult.

With such a design, uGT can store and query almost millions of features of mixed data types for any number of cells if enough storage was given. It can further support more features when features from other omics data are ready to be integrated.

### Uploading data to uGT

The uGT accepts preprocessed data submission via authorized API. In this version, the data were depth-adjusted and log-normalized and followed one consistent format that is ready for uploading. We uploaded 1,093,299 cells to the uGT in total. Every cell is a row with a unique identifier (column “cid”), followed by 43,878 columns of genes expression values and 17 columns of metadata (columns “user_id”, “study_id”, “cell_id”, “organ”, “region”, “subregion”, “seq_tech”, “sample_status”, “donor_id”, “donor_gender”, “donor_age”, “original_name”, “cl_name”, “uhaf_name”, “tissue_type”, “cell_type”, and “marker_gene”) describing dataset-level information and cell-level information.

### ECAUGT: the data access interface of uGT

Based on the Tablestore python SDK, we developed a command line tool “ECAUGT” (pronounced as “e-caught”) to query data from hECA for advanced users to implement *in data* cell sorting. Users can query the cells with the provided query conditions and download the selected data of these cells. For example, the combinatorial query of “all T cell subtypes located in the heart with PTPRC positive and CD3D or CD3E positive” can be written as the following logic expression:

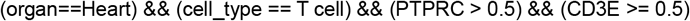

hECA will return all cells that satisfy these conditions in a single downloadable file to users for further analysis. Information about the particular studies of the cells will also be provided to the users. **Table S2** provides the syntax of the logic expressions in ECAUGT.

Function “query_cells()” will query cells with conditions on the columns of metadata and provide a user-friendly interface, with which users can combine multiple conditions into a logical expression in a structured string with logical operators ‘&&’ (for logical operation AND), ‘||’ (for logical operation OR), and ‘!’ (for logical operation NOT). Then “query_cells()” will return the cid list of the queried cells. Function “get_columnsbycell ()” will allow users to download data with this id list. Users can select columns of interest and add gene conditions in this function with the similar interface by “query_cells()”. The “get_columnsbycell ()” can provide downloaded data in two forms: a python list, where each element represents a cell, or a pandas.DataFrame object. User can choose the form they want with the parameter “do_transform”. We also provide the parallel acceleration version with similar interface by “get_columnsbycell_para()”. Function “get_all_rows()” will provide the cid list of all cells in uGT and can be convenient when user require information of the whole hECA. Function “get_column_set()” receives a cid list and will provide all unique values in the selected column of these cells.

For users without much programming background, we provided a lightweight command line tool “Cell_Download” to download data from hECA. Users first query cells in the website interface of hECA and download a cell id list file. Then “Cell_Download” only need one-line command to assign the input and output path and will automatically download all columns of the selected cells in the id list and save the result in four files: a .csv file “metadata.csv” for columns of metadata, a .npz file for sparse expression matrix, and two .csv files for the row names and column names of this matrix. “ECAUGT” is available on https://pypi.org/project/ECAUGT and can be installed with PyPI. Full documentation of ECAUGT could be found at http://eca.xglab.tech/ecaugt/index.html.

### Constructing the unified hierarchical cell annotation framework (uHAF)

To assemble cells into an atlas so that cell annotations from different studies can be aligned, we designed an index and coordinate system, uHAF. The uHAF is a structured framework we designed for the and hierarchical indexing and annotation of organ origin and cell types in hECA. We unified the information of anatomical structures, source organs, and cell types into a unified hierarchical knowledge graph. Users can assign annotations at multiple granularities with uHAF, depending on the quality of the data to be labeled.

We defined two types of entities using a controlled vocabulary, composing two subgraphs in uHAF. Entities in the “macroscopic subgraph” include system, organ, anatomical region and subregion information (see “uHAF macroscopic ontology” in https://github.com/XuegongLab/hECA/tree/main/UHAF). Entities in the “microscopic subgraph” include annotations of cells on their histological types (epithelial tissue, connective tissue, muscle tissue and nerve tissue) and cell types or subtypes determined by molecular features (see “uHAF microscopic ontology” in https://github.com/XuegongLab/hECA/tree/main/UHAF). We defined two types of edges in the uHAF graph, “part of” and “is a”, to represent the hierarchical relations among the entities, and an extra “connect to” type of edge to tag attributes of the entities. For example, there is a “part of” edge from the entity “left ventricle” to the entity “heart”, and there is an “is a” edge from the entity “inhibitory neuron” to the entity “neuron”. If a cell type present in certain organs, there are “part of” connections from cell type nodes to organ nodes, indicating the cell type composition of a macroscopic entity. For example, the entity “T cell” has a “part of” connection with the entity “left ventricle”, as well as “part of” connections to other anatomical units that have T cells in their tissues. We listed all the connection observed in our collected data of hECA v1.0 in “uHAF macro-micro map” (https://github.com/XuegongLab/hECA/tree/main/UHAF).

The entities in the macroscopic and the microscopic subgraph are organized in a hierarchical DAG (directed acyclic graph) structure by manually surveying the canonical human anatomy structure and cell type names from classical medical textbooks including *Junqueira’s Basic Histology: Text & Atlas* (Mescher, 2016), *Histology and Embryology* (in Chinese) (Tang and Zhang, 2013), *Systematic Anatomy* (in Chinese) (Bai and Ying, 2015), *Histology and Embryology* (in Chinese) (Li and Zeng, 2018) as well as several public studies and databases (Franzén *et al*., 2019), followed by confirmation and refinement from medical experts. We then organized the macroscopic and the microscopic subgraphs into ontologies by the protégé tool (https://protege.stanford.edu/).

The microscopic entities are attached with attributes “marker reference” consisting of marker genes by the “connect to” edges (see “uHAF marker reference” in https://github.com/XuegongLab/hECA/tree/main/UHAF). We manually collected these marker genes that are often used in articles. For cell types whose marker genes were not given in the original studies, we surveyed for markers from multiple sources including PanglaoDB (https://panglaodb.se/) (Franzén *et al*., 2019), the Human Protein Atlas (https://www.proteinatlas.org/)(Uhlén *et al*., 2015), and CellMarker (http://biocc.hrbmu.edu.cn/CellMarker/)(Zhang *et al*., 2018) to replenish the marker references. Such processes were implemented iteratively to curate the final marker references. The references will be continuously updated along with the release of new versions of hECA.

We provided the uHAF related files at https://github.com/XuegongLab/hECA.

### Cell identity assignment

We assigned an identity label from the uHAF to every single cell collected. Each cell in hECA is annotated with two entities of the uHAF, one macroscopic and one microscopic. The annotation can be of different levels in the two hierarchies, depending on the information provided by the original data and the specificity of the marker genes.

#### uHAF name assignment

For each Seurat cluster, we identified the cluster-specific differentially expressed genes (DEGs) by FindAllMarkers function. We referred to the marker reference to determine the cell type labels, and use the top ranked DEGs to further annotate the subtypes. We first determined the most general labels among the four tissue types (epithelial tissue, connective tissue, muscle tissue, nerve tissue), and then chose the deepest child cell type on which markers can be used to support the cell type assignment in the uHAF. In this way, we annotate each cluster “organ-tissue_type-cell_type-markers”, indicating the macroscopic and microscopic level of the cluster. For cells that cannot be annotated based on available information, we named them as “Unclassified”. This label produced from uHAF is called “uHAF_name”. **Table S3** listed the entity combinations that have been used in annotating the existing data in the current version of hECA. Users can use the uHAF to annotate their query cells in the same way.

#### Mapping uHAF names to Cell Ontology terms

We downloaded the basic Cell Ontology (Bard et al., 2005; Diehl et al., 2016) terms from CL website (Cell Ontology - Summary | NCBO BioPortal (http://bioontology.org)), retained “Preferred Label”, “Definitions” and “Parents” (Table), and used the “Preferred Label” for CL term assignment. We converted the “uHAF_name” to “cl_name” by a combined strategy: We preferably used the Cell Ontology terms with the exact matching of the whole string of “cell_type”. For the “cell_type” that did not appear in the Cell Ontology terms, we further searched their parent “cell_type” in our uHAF until the Cell Ontology term is matched completely. For the remaining “cell_type”s, we manually determined the most similar Cell Ontology terms by ontologyIndex R package (Greene *et al*., 2017). If no term was found after these steps, we labeled them as “none” (see **Table S3**).

### Generation of quantitative portraits

We designed a portraiture system as a systematic way to characterize the full properties of biological entities of all levels in hECA. There are three major types of biological entities in hECA: organs (including sub-organs), cell types (including subtypes) and genes. A full quantitative portrait of a biological entity is its holographic picture of the entity at anatomical, cellular and molecular levels. However, both the quality and quantity of the currently available data in hECA are far from constructing such full portraits. Therefore, the quantitative portraits in hECA v1.0 only illustrated the idea of the portraiture system using the available information. They reflect more about the characteristics of the collected data of and related to each entity, rather than about the biological truth of the entity.

Organ portraits: a portrait of an organ is composed of 3 major parts: the cell composition viewer, the cell embedding viewer, and the organ hierarchy viewer. The cell composition viewer shows the counts and fractions of cell types observed in one organ’s datasets. It is notable that statistics in the organ portraits only reflect the counts/fractions of the collected cells, not the true counts/percentages of cell types in an organ. The embedding viewer visualizes cells of an organ with a 2-dimensional scattergram (UMAP/PCA/DensMAP for users to choose). This viewer supports coloring embedded cells by their cell types, sequencing technologies, original studies, and any given gene’s expression level. The organ hierarchy viewer shows the position of the organ in the uHAF macroscopic annotation system.

Cell type portraits: The cell type portrait depicts cells belong to the same cell types/subtypes across all organs, and is composed of 4 major parts: cell distribution, marker genes, 2D visualization and taxonomy relationship with other cell types. The cell distribution part describes the relationship of this cell type with organs, with bar plots showing the organ origin of cells in numbers and proportions. The marker gene part provides a table with genes highly expressed in this cell type, which were defined by comparing gene expression level with all other cell types using Seurat v3.2. We filtered out genes with adjusted p-value larger than 0.05 or expressed in fewer than 25% cells in this cell type, and showed top 50 genes with highest log fold-changes. In 2D visualization part, we plotted an interactive scatter plot showing the distribution and landscape of cells in this cell type. Embedded cells can be colored with their organs, sequencing technologies, original studies, and any given gene’s expression level. Like organ portraits, we also showed the cell type’s hierarchical relationship with other uHAF cell types.

Gene portraits: The portrait of a gene is composed of 2 major parts: basic gene information and gene expression distribution. In the basic gene information part, for each gene, we collected the full name of gene, the position where the gene is on the genome, commonly used aliases of the gene, and description that introduce the basic function of the gene. The “known as marker of” section denote cell types that highly express this gene, which is calculated by comparing the expression level of the gene in a cell type with it in other cells. For the gene expression distribution part, we first performed data normalization of all cells in uGT using function NormalizeData in Seurat v3.2. For each gene, we present its distribution in an organ or in a cell type by drawing a ridge plot. The ridge plot is fitted by expression value of the gene in the organ or cell type, while zero-value are truncated before fitting. The median expression level and non-zero percentage are also provided on the ridge plot.

### The hECA website

We provided two portals for users to access hECA. One is a computer programing portal for users to access the data and do *in data* cell experiment using the ECAUGT package. The portal is at https://pypi.org/project/ECAUGT/. It is powerful but requires users to be comfortable with some programming skills. The other portal is a website at http://eca.xglab.tech/ with graphic user interface (GUI) that enables both browsing hECA at all levels and searching the data for *in data* cell experiments. ECAUGT can also be accessed from the website portal.

The interactive functions of the hECA website (http://eca.xglab.tech/) are divided into four parts: “Cell sorting”, “uHAF cells”, “uHAF organs” and “gene portraits”, plus a link to the “ECAUGT” portal. Users can browse these functions anonymously, but signing in is needed to get the full service.

“Cell sorting” is the graphical interface for *in data* cell sorting in hECA v1.0. It supports flexible multi-step cell selection with all kinds of filters regarding to cell features (gene, cell type in uHAF, organ in uHAF and other metadata). Filters can be combined with basic logic operators (AND, OR, NOT) to form complex logic expressions. Users can have a quick view of the selected data with real-time statistical analysis and visualization of the organ origin and cell type composition, and can adjust the sorting criteria accordingly if necessary. For more in-depth analysis, we provide the organ-wise cell type composition and gene expressions across cell types or organs and pseudo-FACS visualization of expression correlation between any two genes. Cell sorting processes can be saved to users’ collections for future reference.

After users selected their interested cell groups, a cid list can be downloaded for further data query with ECAUGT. Examples of *in data* cell sorting and vignettes are provided in the home page of hECA website.

Cell types and organs are organized in uHAF DAG in hECA. The “uHAF cells” entry provides an interactive tree visualization of the cell types’ hierarchical relationships, which is the microscopic subgraph of uHAF. The “uHAF organ” entry provides the view of the macroscopic subgraph of uHAF. Each cell type or organ is assigned with a unique uHAF ID with a brief description. We provide portraits for cell types with data available in the current version.

Users can click “view details” to check the cell type portraits which include information of original organs, marker genes and embedding view of the cell types. The plots can be colored by the organs, expression level of selected gene, sequencing platform or the original study.

The organ portraits provide information of cell type composition (as reflected by the current data), similar embedding views, anatomy relationships and position in the uHAF.

The “gene portraits” entry allows users to select any particular gene and visualize the distribution of the gene in all organs and cell types (as reflected by the currently available data). The basic information includes the distribution of non-zero expression values in the organs and cell types, and the proportion of non-zero values (%Expr). Users should keep in mind the fact that the current scRNA-seq data are quite noisy and suffer from dropout events when using the information. The gene portraits also provide basic information of the gene collected from public databases and links to the corresponding pages at Genecard, NCBI, Ensembl and Wikigenes.

## SUPPLENMENTAL INFORMATION

**Table S1**. Source datasets of hECA v1.0. Related to **Table 1**.

**Table S2**. Syntax of the logic expressions of ECAUGT in uGT. Related to **Figure 1**.

**Table S3**. uHAF annotation produces uHAF_name. Related to **Figure 1**.

**Table S4**. Details of case study 1: Negative markers for T cells. Related to **Figure 2**.

**Table S5**. Details of case study 1: Positive markers for T cells. Related to **Figure 2**.

**Table S6**. Details of case study 2: Organ and cell type table of CD19+ cells. Related to

**Figure 3**. The number of CD19 expressed cells from each cell type and each organ. Red and blue highlights were treatment effect and side effect respectively.

**Table S7**. Details of case study 2: Organ and cell type table of CD22+ cells. Related to

**Figure 3**. The number of CD22 expressed cells from each cell type and each organ. Red and blue highlights were treatment effect and side effect respectively.

**Data S1**. Organ cellular composition. Related to **Table 1**. Each page shown one organ and its cell type composition. The cell number and percentage were shown for each cell type. Database query was performed using the function “get_column_set()” of package ECAUGT to extract the cells for each cell type, and each cell type per organ was counted by “express.treemap()” function in “ploty” python package.

**Figure S1**. Filtering candidate T cell subpopulations. Related to **Figure 2**. (A) A UMAP showing the within-organ clustering results. (B) The general T cell markers’ expressions (CD3D, CD3E, CD3G). (C) Per-cluster marker gene expressions.

**Figure S2**. CD4/CD8 T cell population definition. Related to **Figure 2**. (A) A heatmap showing the hierarchical clustering results based on the CD4/CD8A/CD8B genes. Other T cell signature genes are also listed on the heatmap. (B) Split view of CD4 positive T cells, CD8 positive T cells, and double-negative T cells (Note that sequencing dropouts caused some falsely recognized double-negative T cells).

**Figure S3**. Cell type and organ distribution of CD19 expressed cells. Related to **Figure 3**. The number of CD19 expressed cells from different organs (A) and cell types (B). Visualization of CD19 expressed cells in UMAP labeling organs (C) and cell types (D).

**Figure S4**. Cell type and organ distribution of CD22 expressed cells. Related to **Figure 3**. The number of CD22 expressed cells from different organs (A) and cell types (B). Visualization of CD22 expressed cells in UMAP labeling organs (C) and cell types (D).

**Figure S5**. Example gene portrait page of PTPRC. Related to **Figure 1**. Basic information about PTPRC and distribution of expression level grouped by organs are shown in the “About the Gene” and “Gene Expression Profiling” panel, respectively. Users can select which gene to visualize and group by either organ or cell types in the “Select a Gene” panel.

**Figure S6**. Example cell type portrait page of Fibroblast. Related to **Figure 1**. A brief description of Fibroblast is provided under the title. “Original organs” section shows the number and proportion of fibroblast cells in each organ. “Marker genes” section lists the differentially expressed genes in fibroblast and their characteristic expression ranges. “Embedding viewer” section visualizes the distribution of cells on 2D DensMAP space, colored by organs. It also provides 2D PCA and UMAP visualization of cells colored by sequencing technology, data source or expression level of certain gene. “Position on uHAF” marks the position of Fibroblast on the unified hierarchical annotation framework (uHAF).

**Figure S7**. Example organ portrait page of Brain. Related to **Figure 1**. A brief description of Brain is provided under the title. “Cell type composition” section shows the number and proportion of brain cells in each cell type. “Embedding viewer” section visualizes the distribution of cells on 2D UMAP space, colored by cell types. It also provides 2D PCA and DensMAP visualization of cells colored by sequencing technology, data source or expression level of certain gene. “Anatomy relationship” shows the brain belongs to the nervous system and is related to eye and skin. “Position on uHAF” marks the position of Brain on the unified hierarchical annotation framework (uHAF).

