## Supplementary Figures and Tables for "hECA: the cell-centric assembly of a cell atlas": Supp figure and data_1102.pdf

Supplementary figures and data

Figure S1. Filtering candidate T cell subpopulations. Related to Figure 2.

Figure S2. CD4/CD8 T cell population definition. Related to Figure 2.

Figure S3. Cell type and organ distribution of CD19 expressed cells. Related to Figure 3.

Figure S4. Cell type and organ distribution of CD22 expressed cells. Related to Figure 3.

Figure S5. Example gene portrait page of PTPRC. Related to Figure 1.

Figure S6. Example cell type portrait page of Fibroblast. Related to Figure 1.

Figure S7. Example organ portrait page of Brain. Related to Figure 1.

Data S1. Organ cellular composition. Related to Table 1.

Figure S1. Filtering candidate T cell subpopulations. Related to Figure 2.

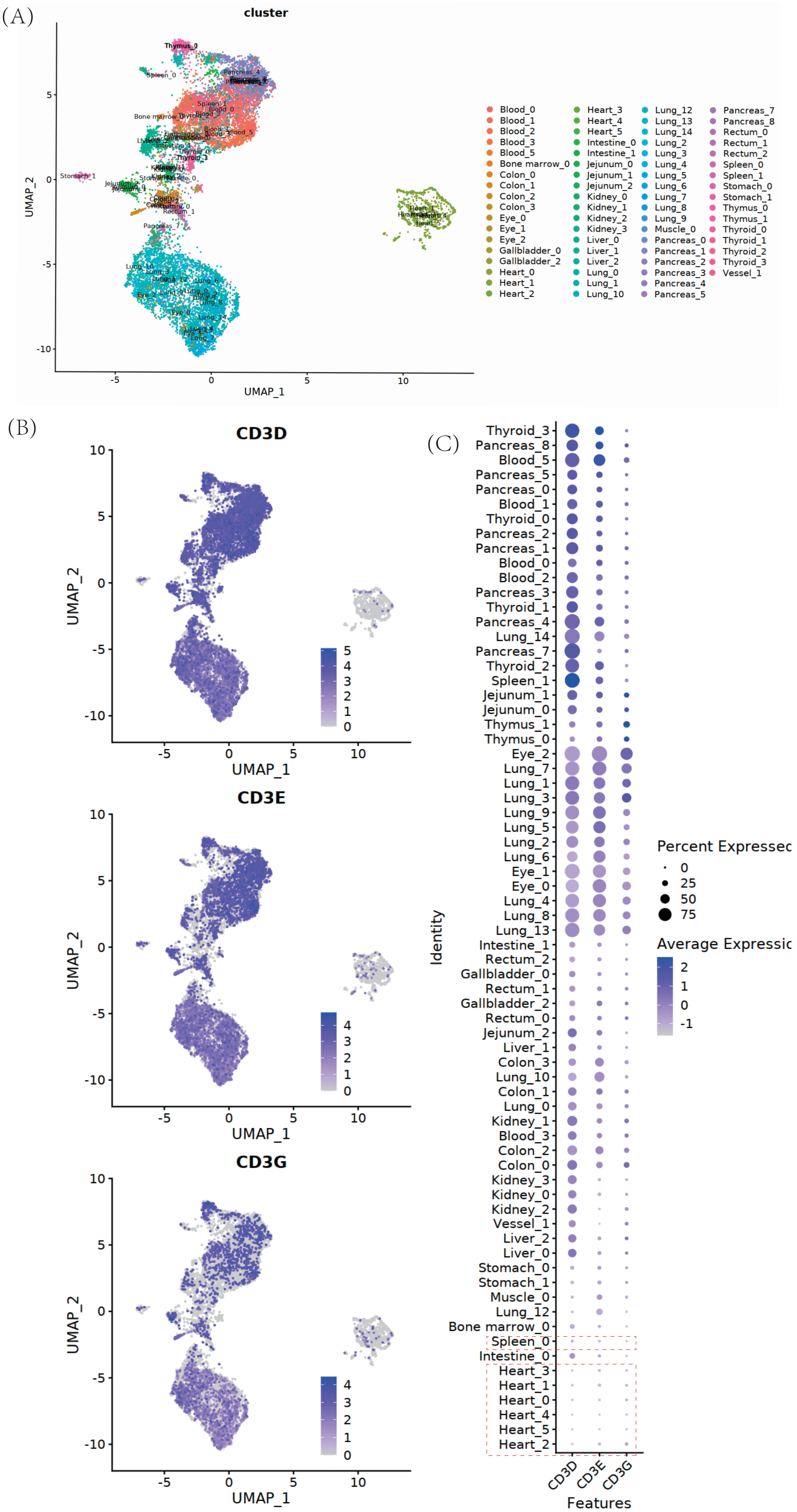

### Figure S2. CD4/CD8 T cell population definition. Related to Figure 2.

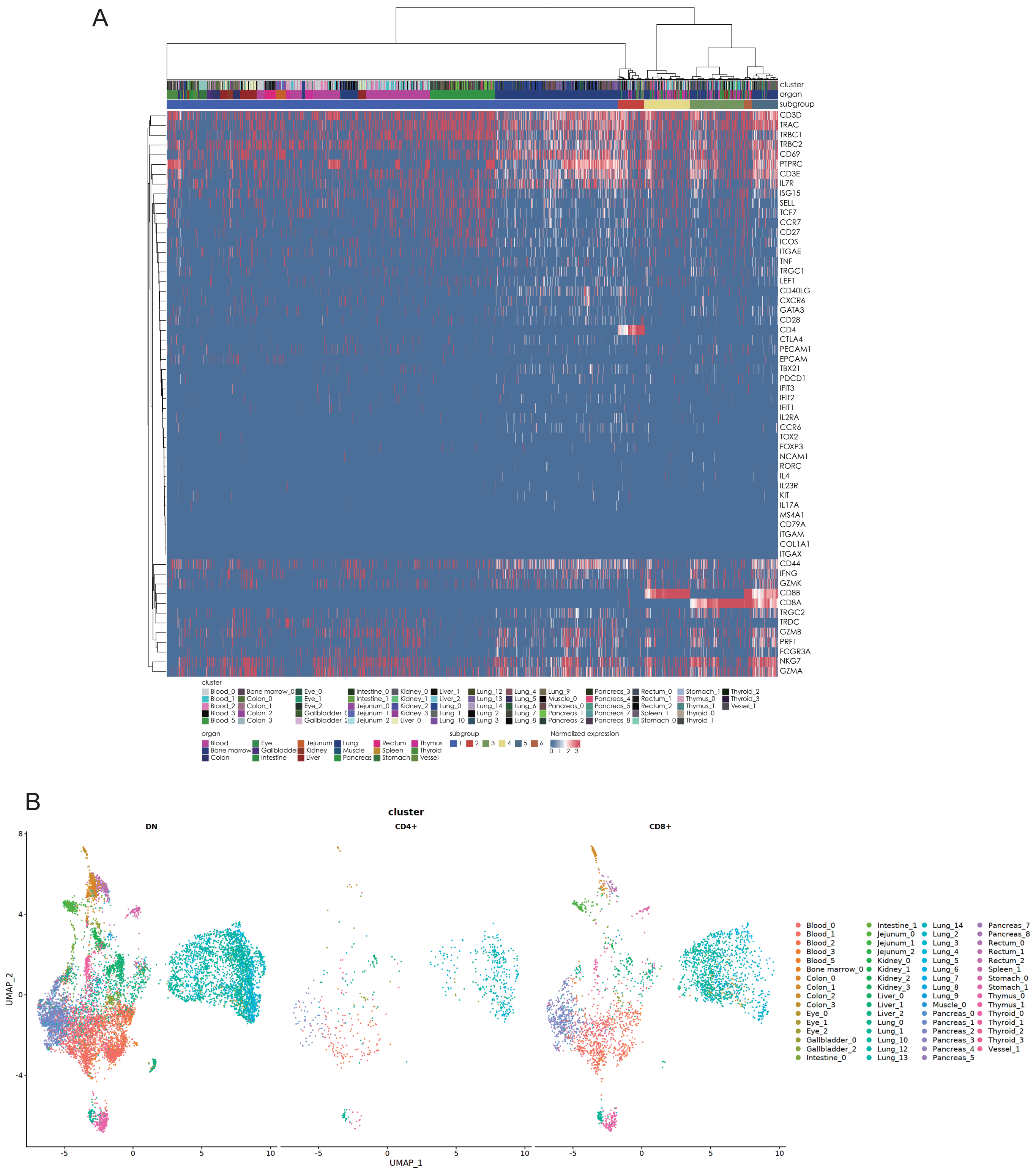

Figure S3. Cell type and organ distribution of CD19 expressed cells. Related to Figure 3.

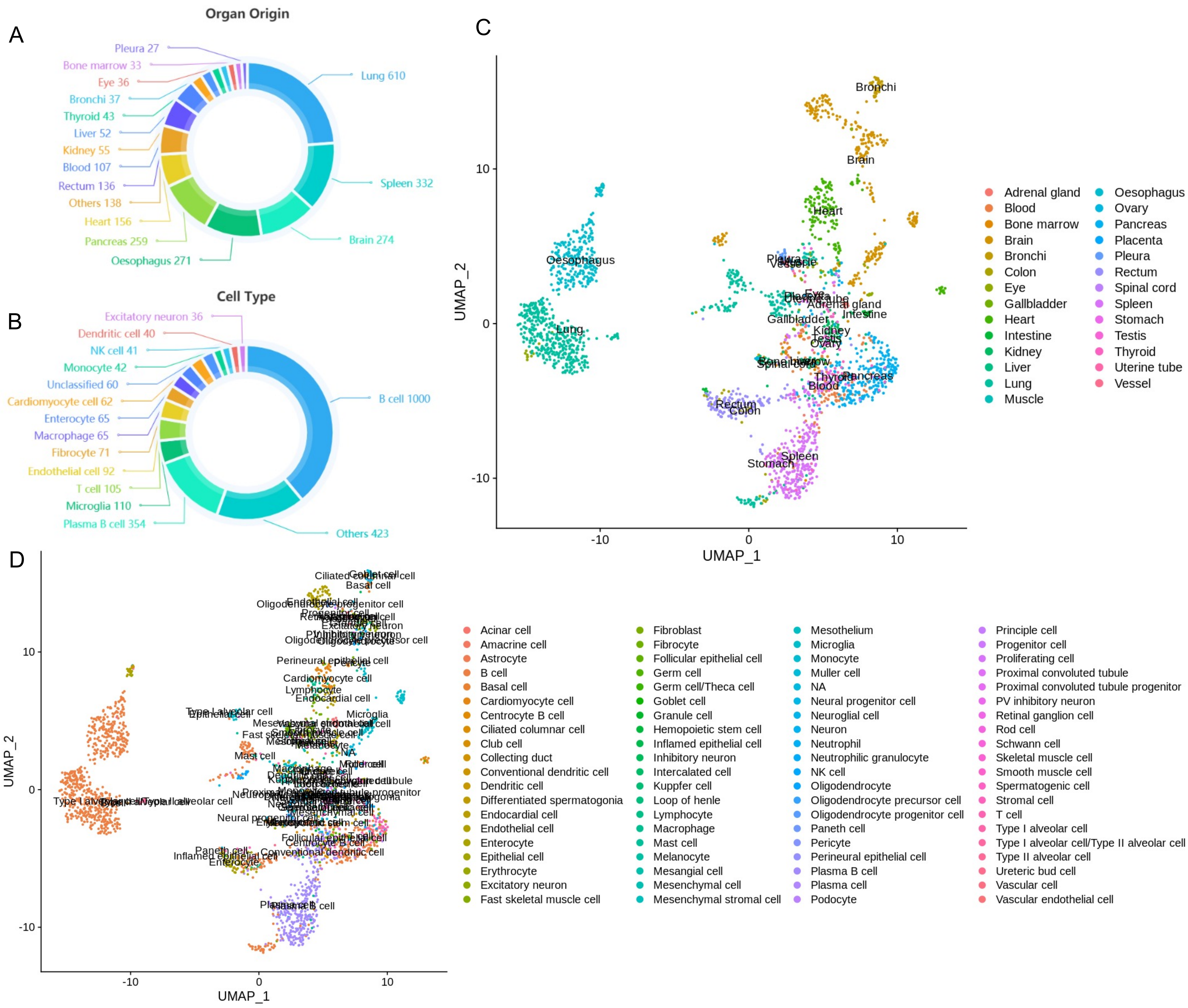

Figure S4. Cell type and organ distribution of CD22 expressed cells. Related to Figure 3.

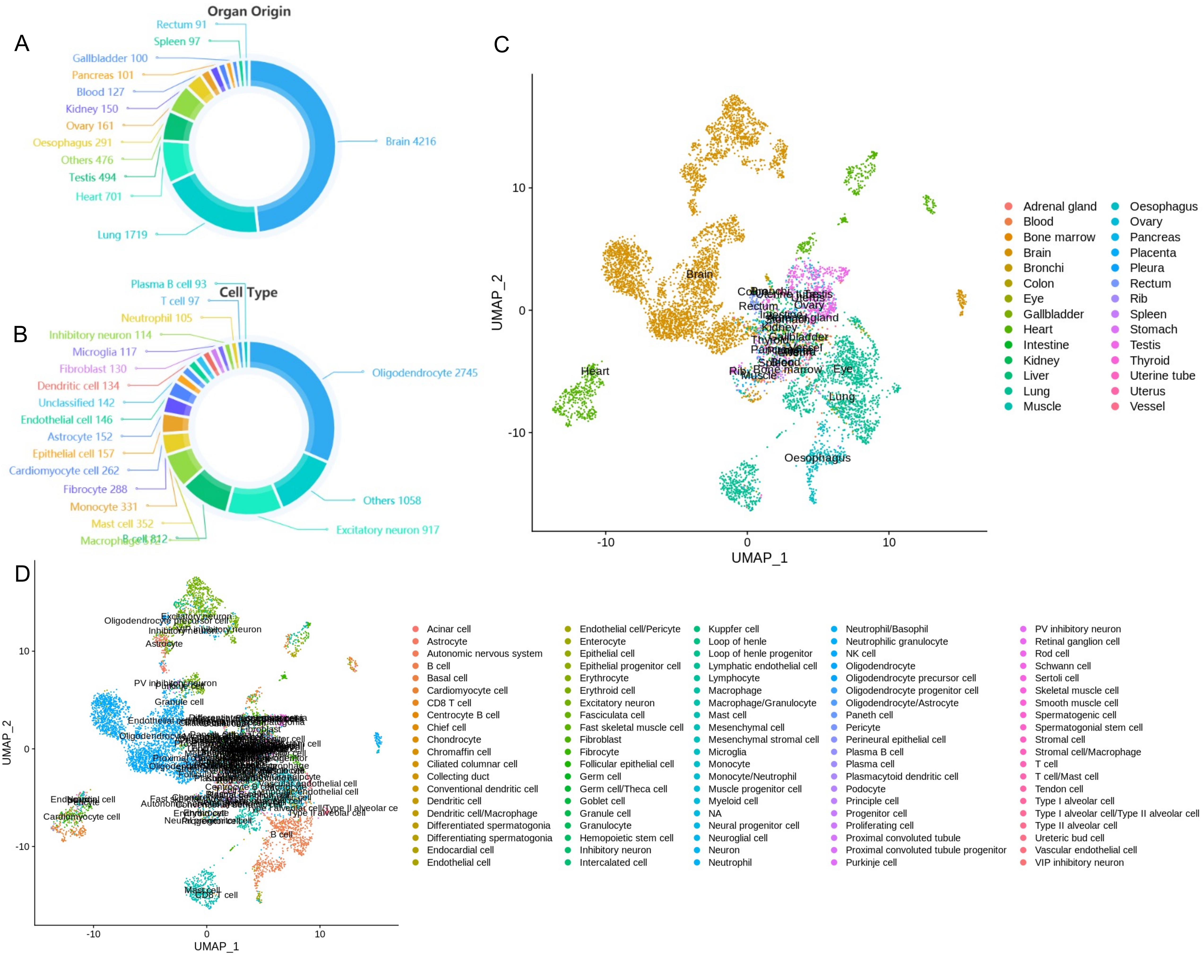

Figure S5. Example gene portrait page of PTPRC. Related to Figure 1.

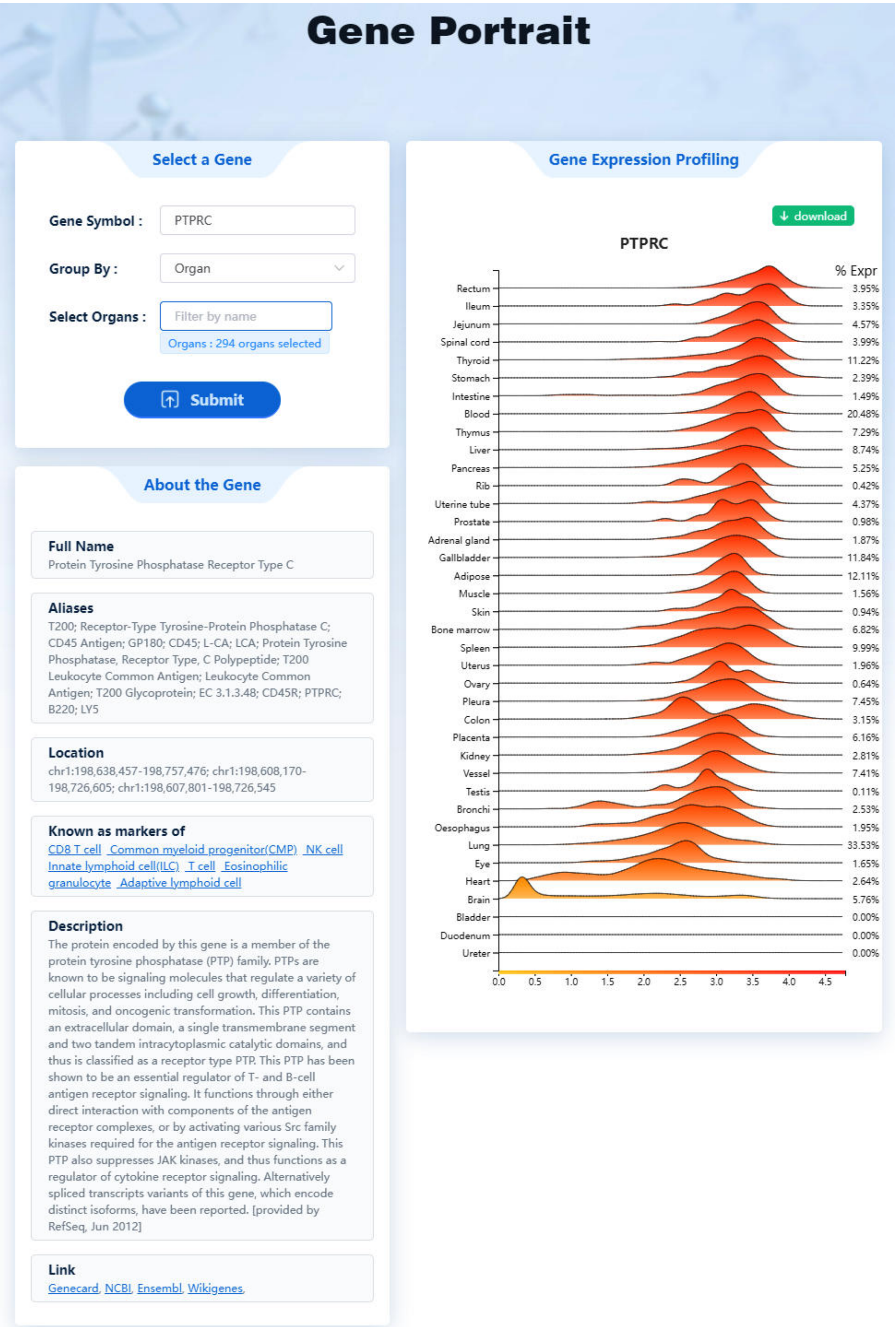

Figure S6. Example cell type portrait page of Fibroblast. Related to Figure 1.

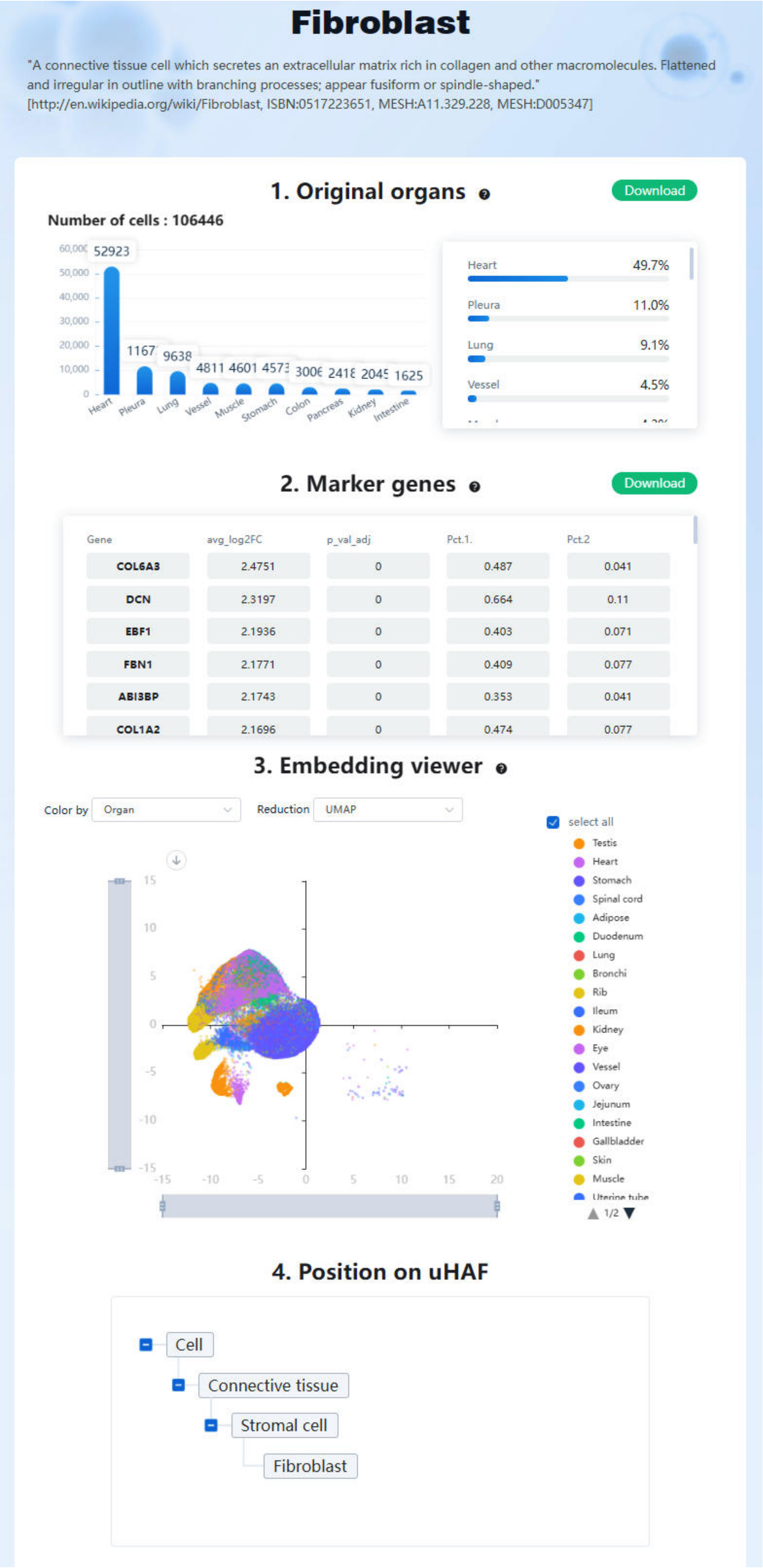

Figure S7. Example organ portrait page of Brain. Related to Figure 1.

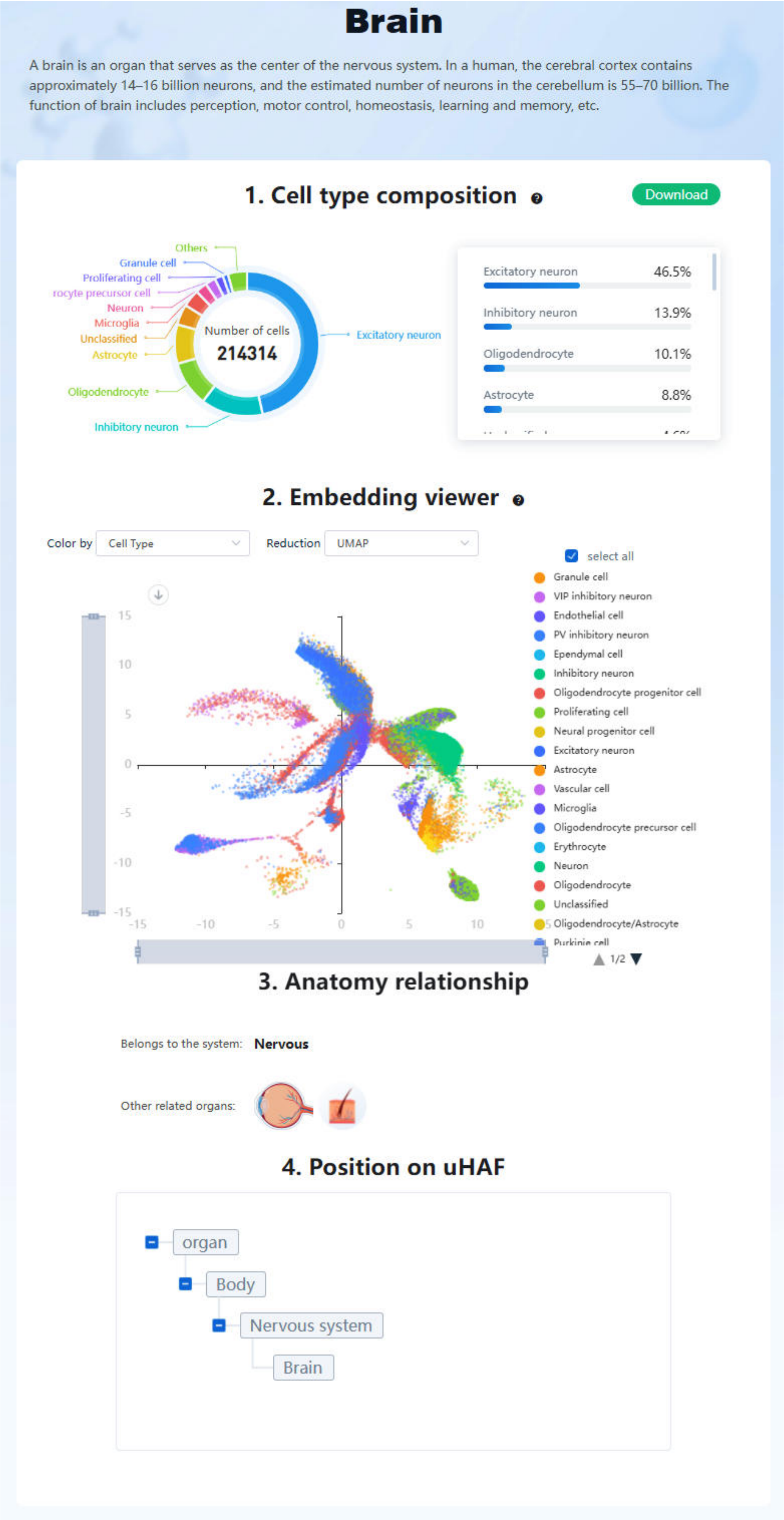

### Data S1. Organ cellular composition. Related to Table 1.

Organ:Ovary   Cell number:6549

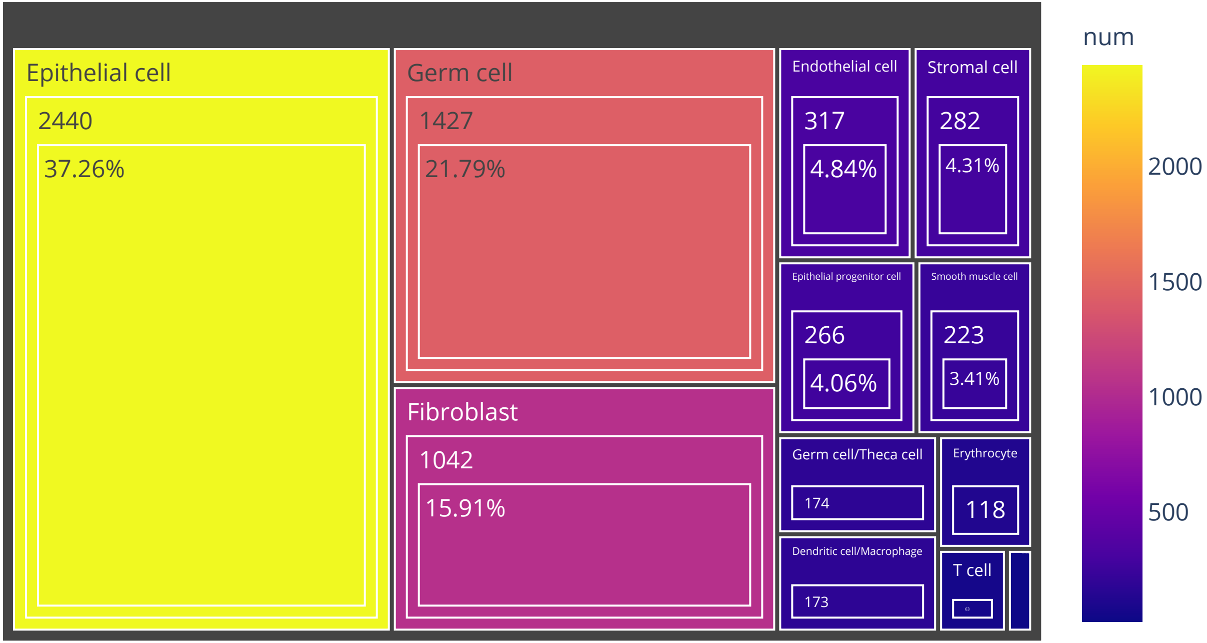

Organ:Bladder   Cell number:3980

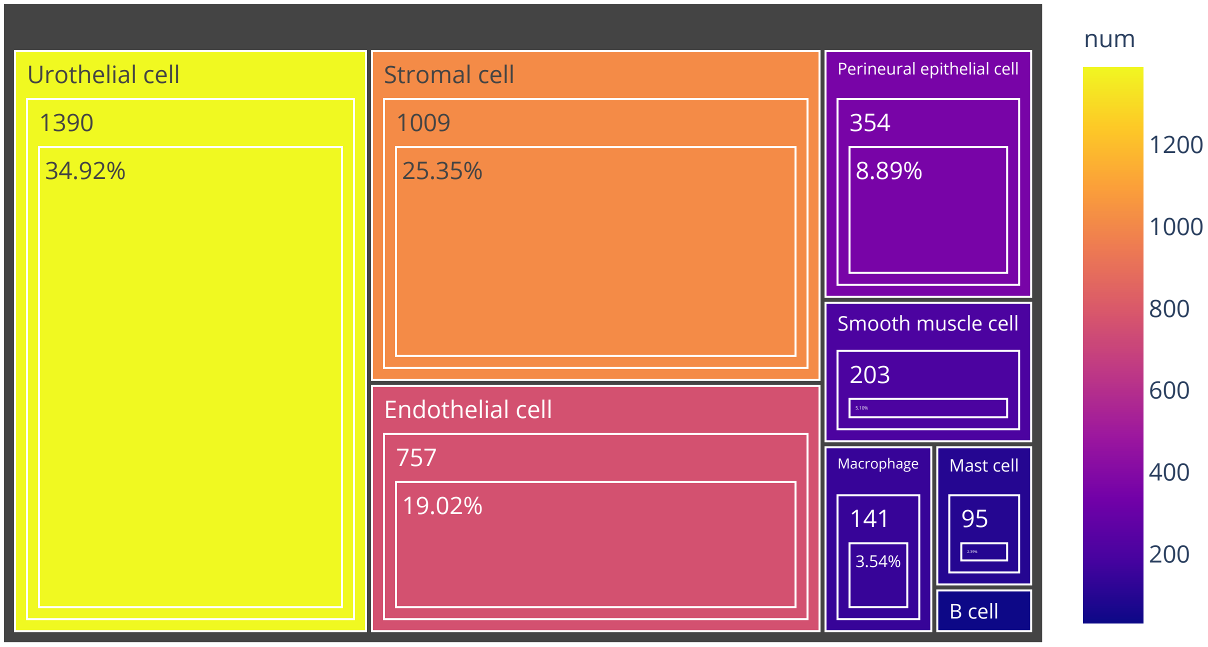

# Organ:Duodenum   Cell number:3743

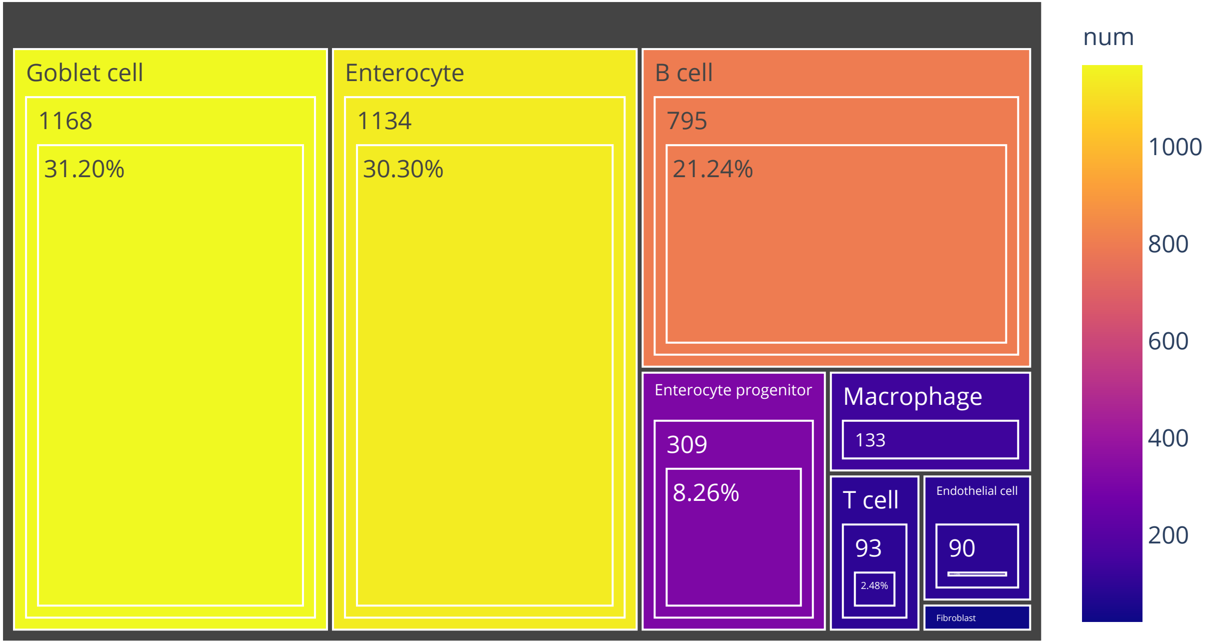

Organ:Thymus   Cell number:4516

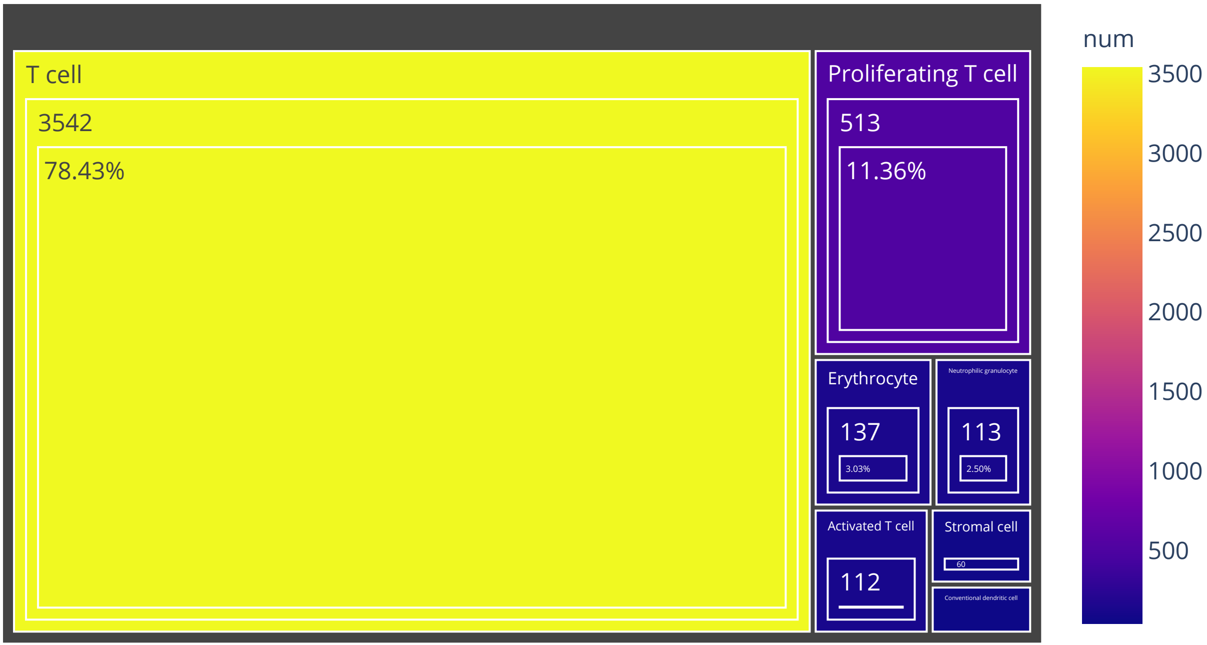

## Organ:Jejunum   Cell number:4198

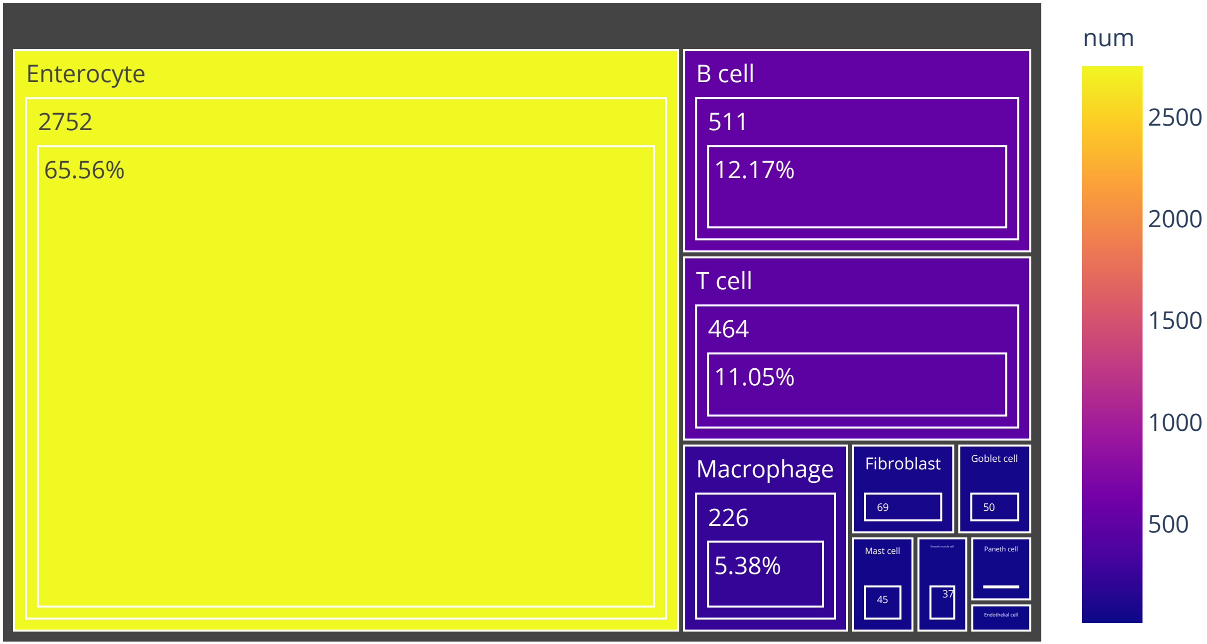

# Organ:Testis   Cell number:12769

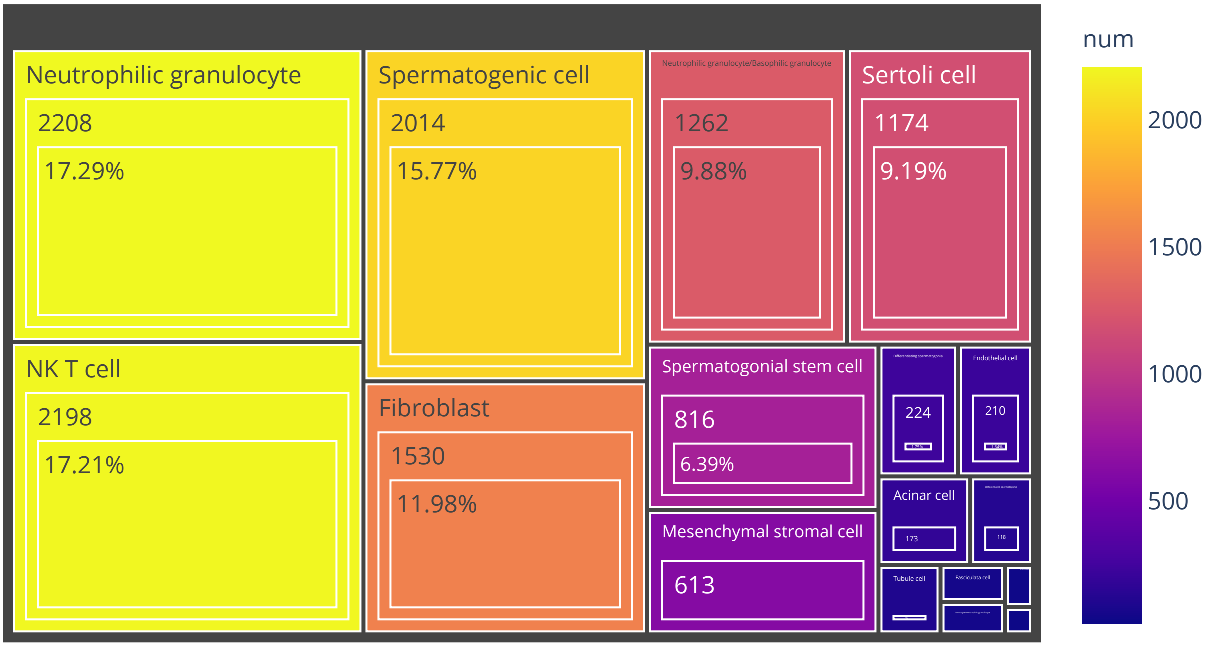

# Organ:Skin   Cell number:6618

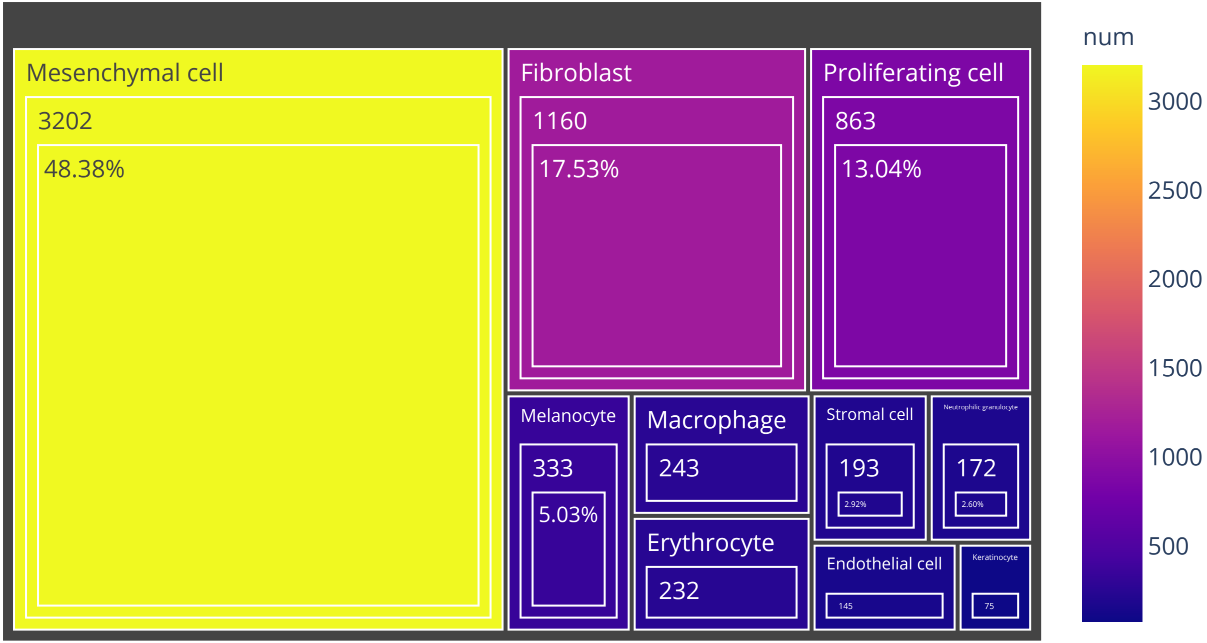

# Organ:Brain   Cell number:214314

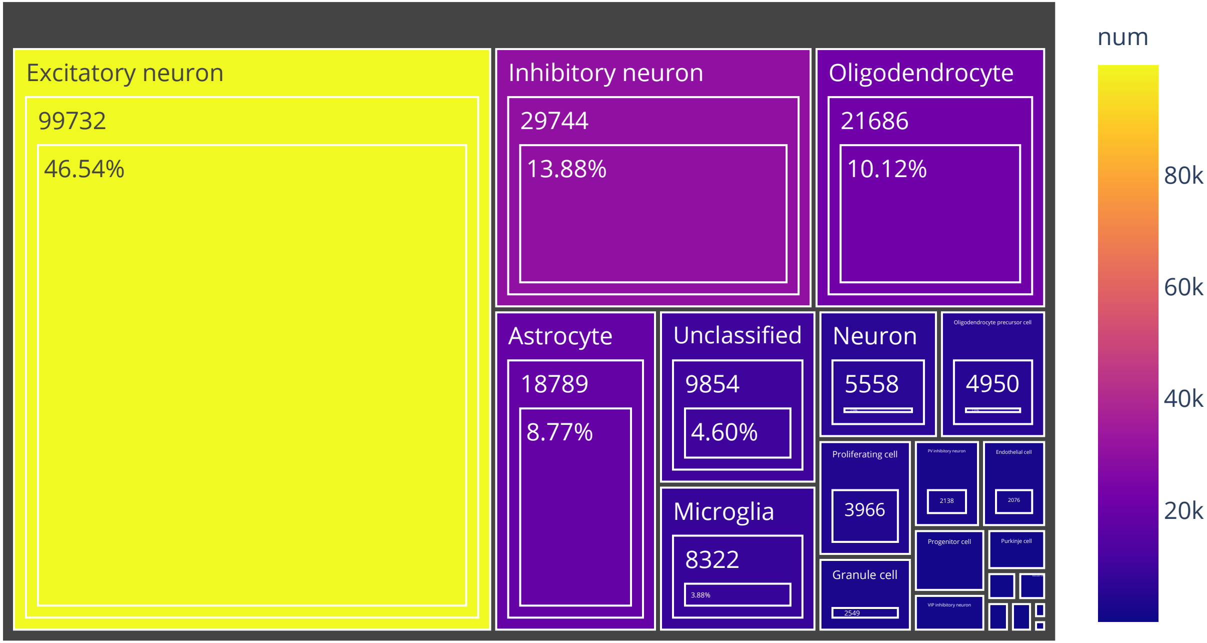

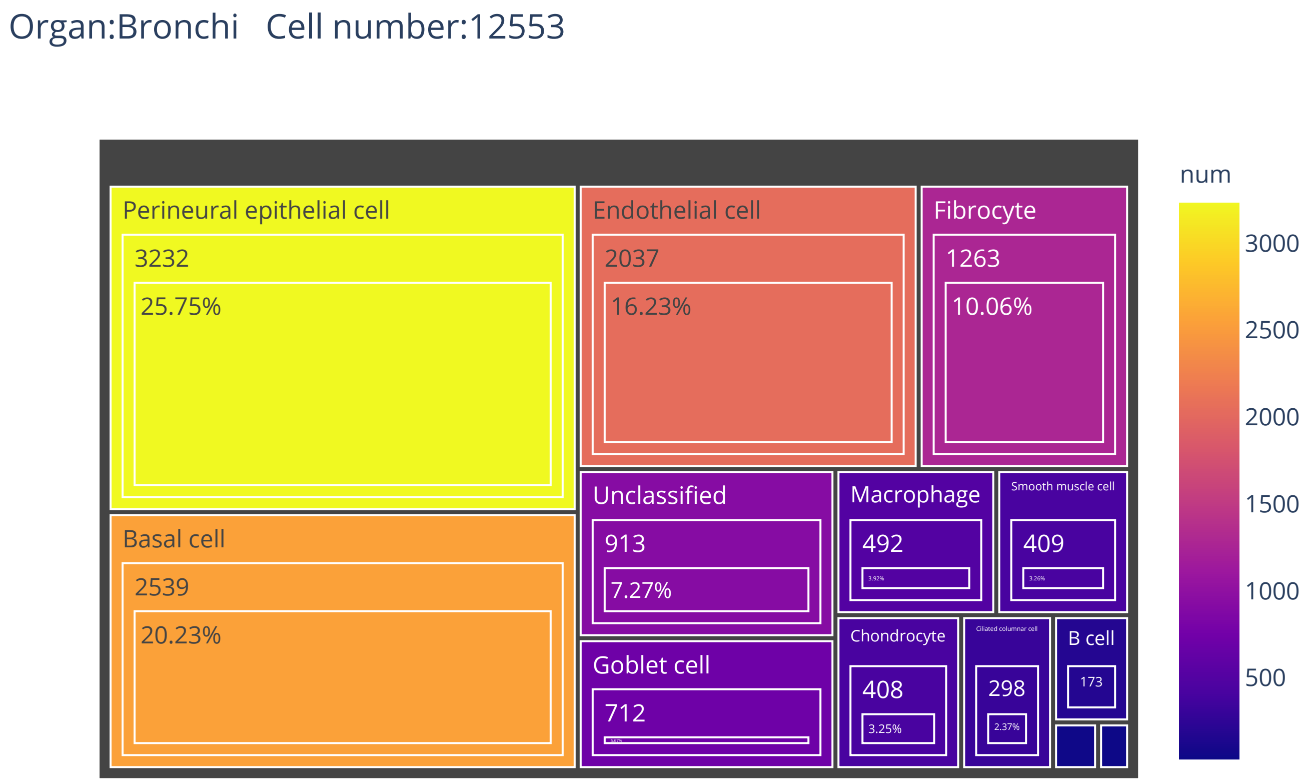
Organ:Bronchi   Cell number:12553
Perineural epithelial cell
3232
25.75%
Endothelial cell
2037
16.23%
Fibrocyte
1263
10.06%
Basal cell
2539
20.23%
Unclassified
913
7.27%
Macrophage
492
3.92%
Smooth muscle cell
409
3.26%
Goblet cell
712
5.67%
Chondrocyte
408
3.25%
Ciliated columnar cell
298
2.37%
B cell
173
num
3000
2500
2000
1500
1000
500

Organ:Placenta   Cell number:9926

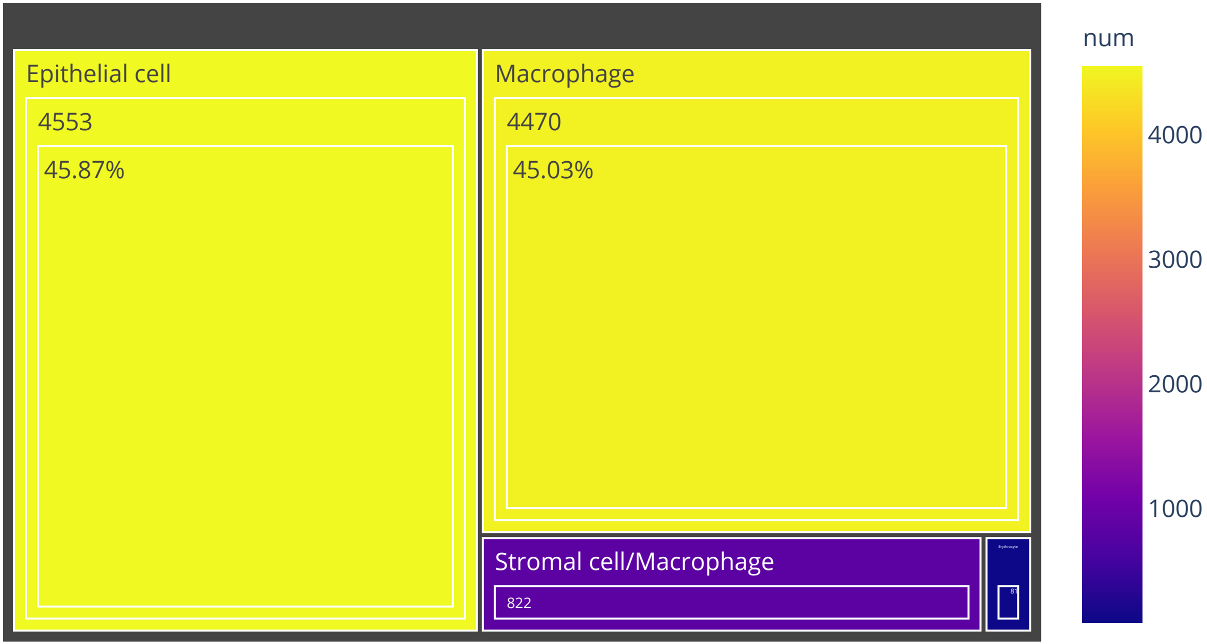

Organ:Blood Cell number:29514

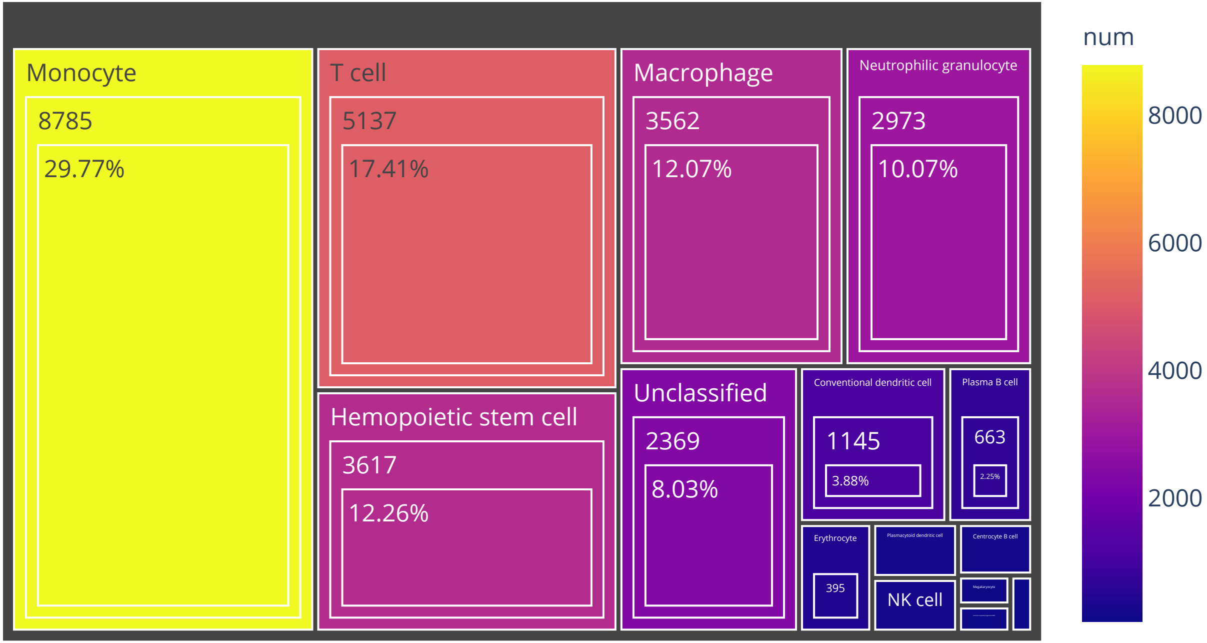

Organ:Prostate   Cell number:2445

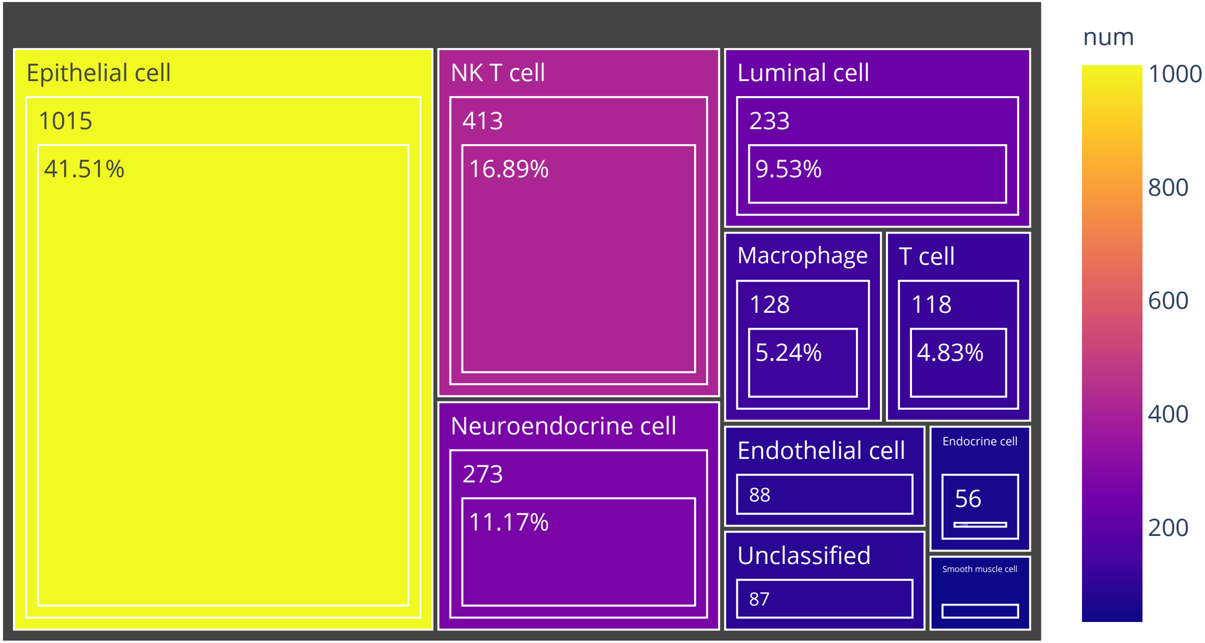

Organ:Lung   Cell number:90521

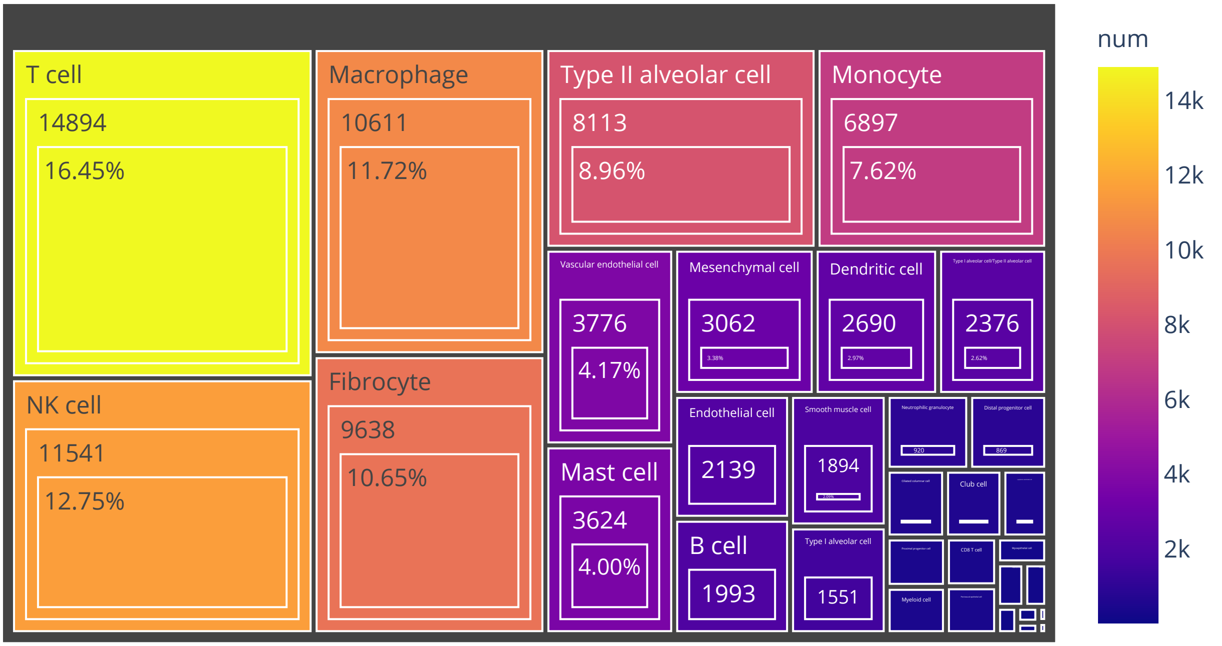

# Organ:Uterine tube   Cell number:6496

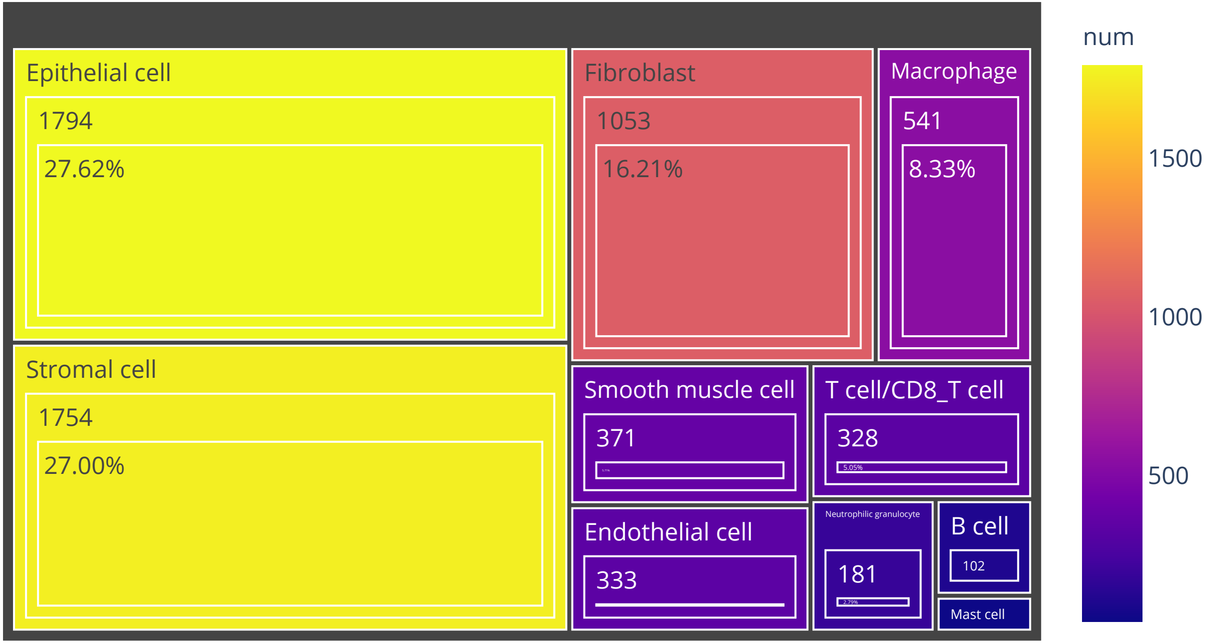

Organ:Adrenal gland   Cell number:15065

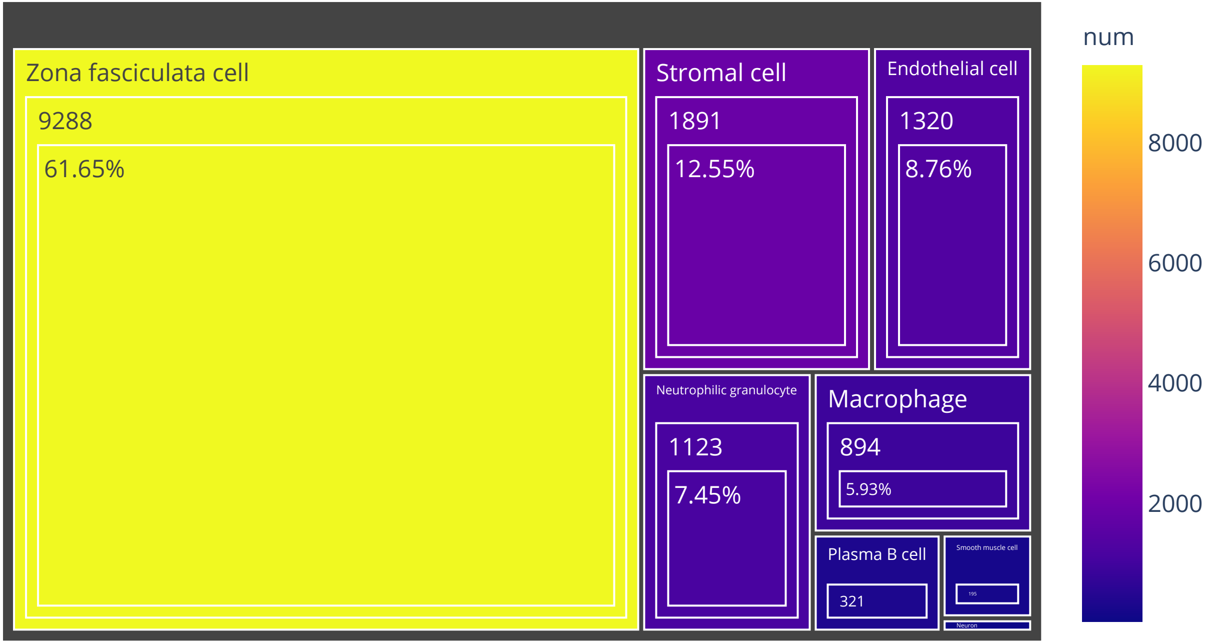

### Organ:Rectum Cell number:5718

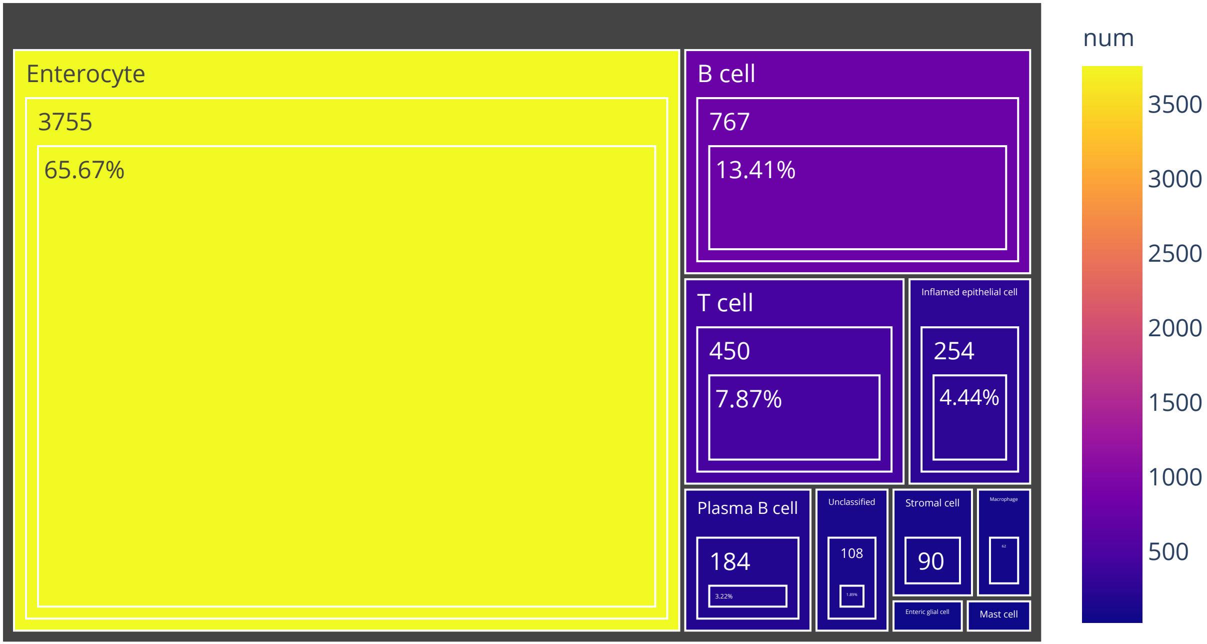

### Organ:Stomach Cell number:22187

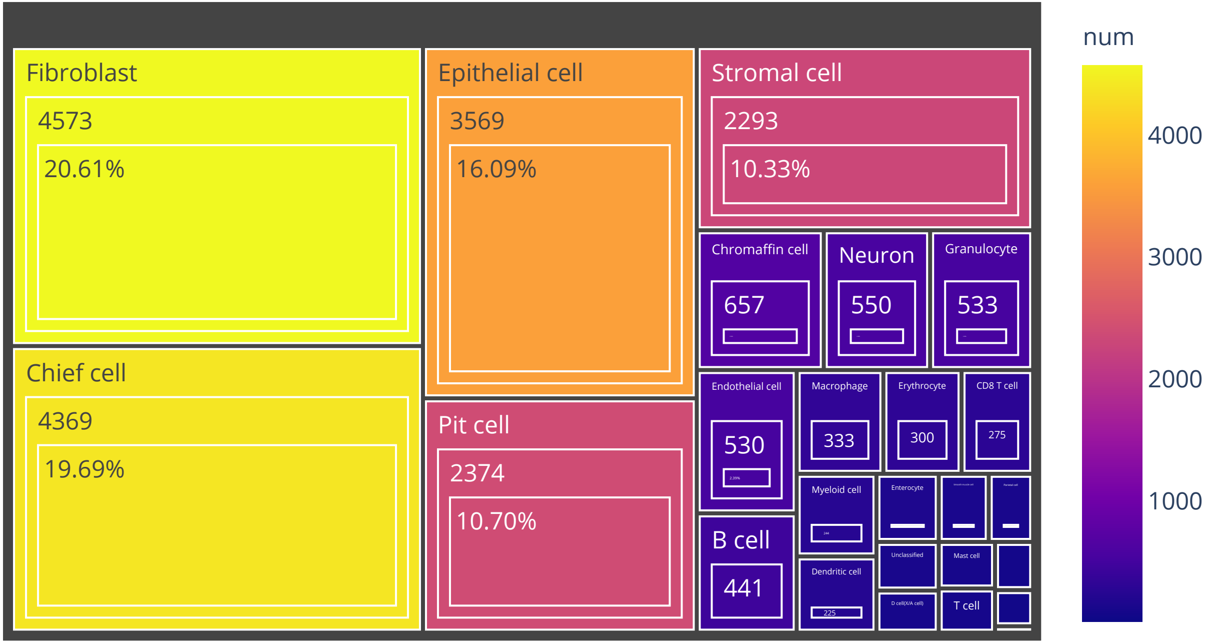

Organ:Ileum   Cell number:3132

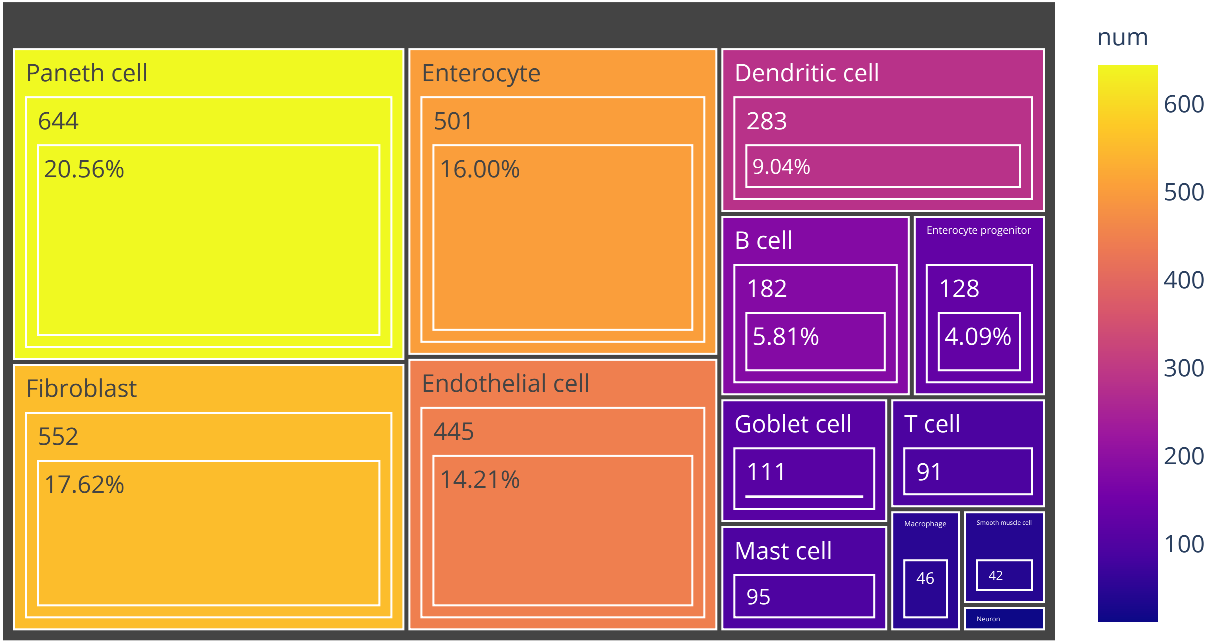

Organ:Spleen   Cell number:15806

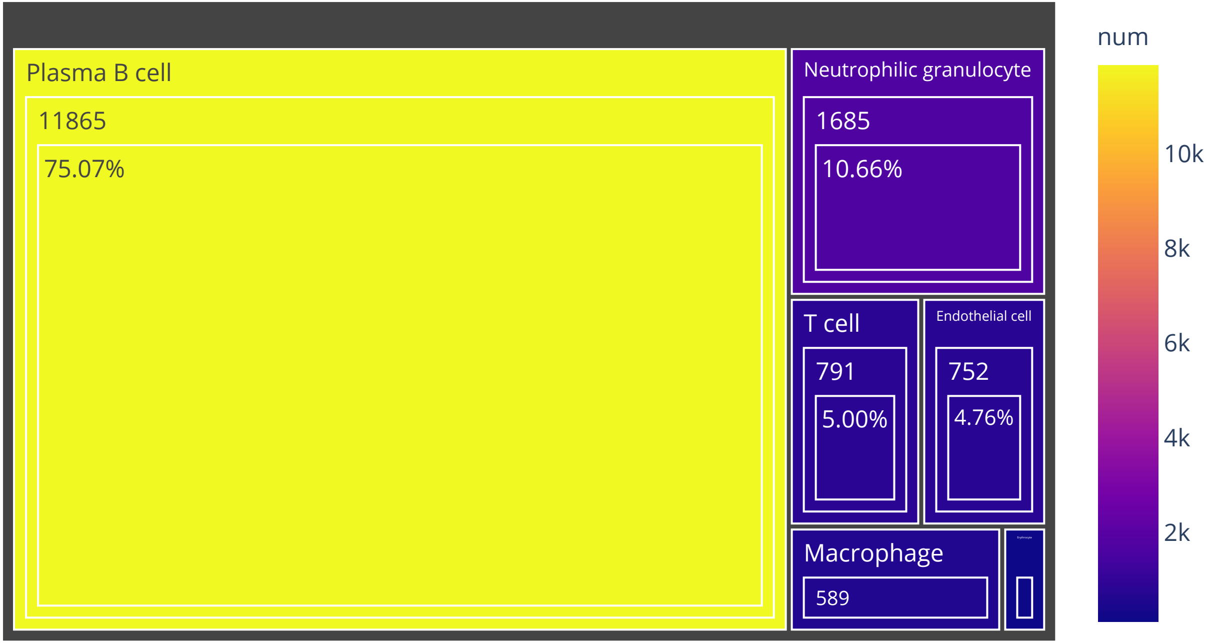

# Organ:Eye   Cell number:47275

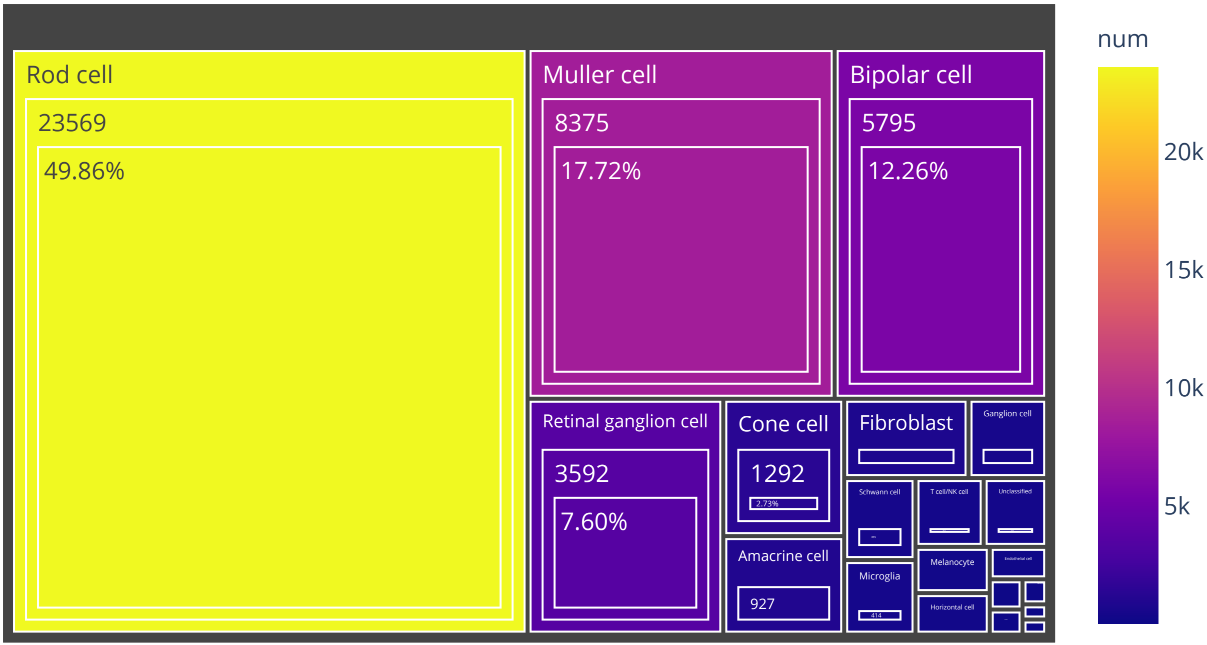

# Organ:Ureter   Cell number:2160

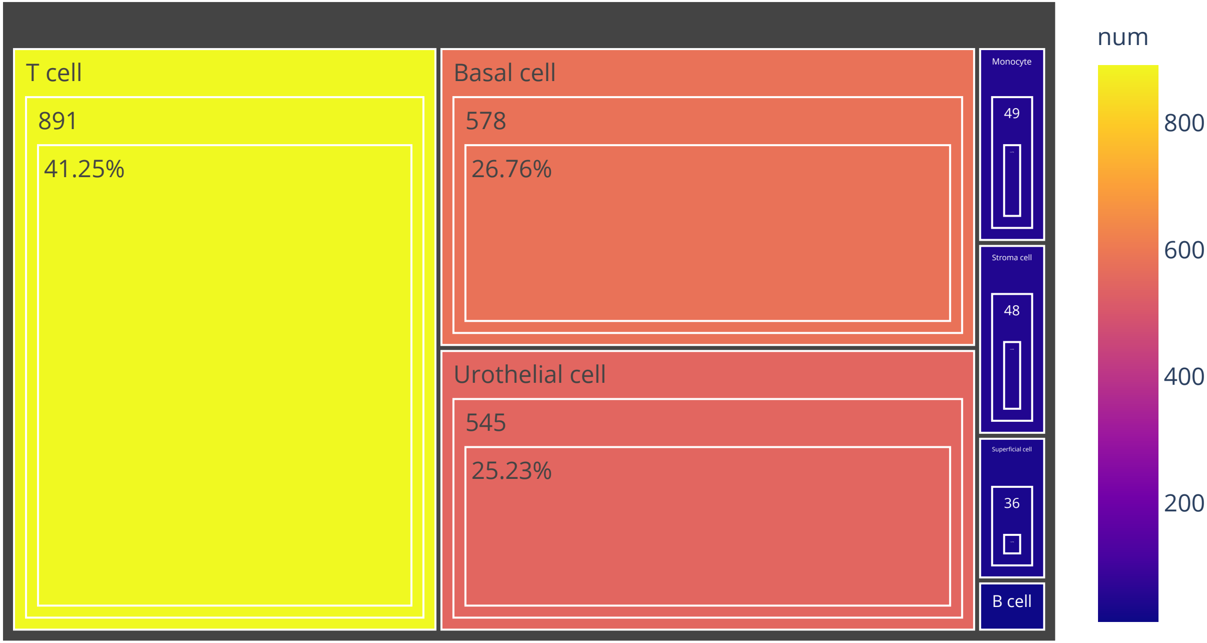

Organ:Adipose   Cell number:1362

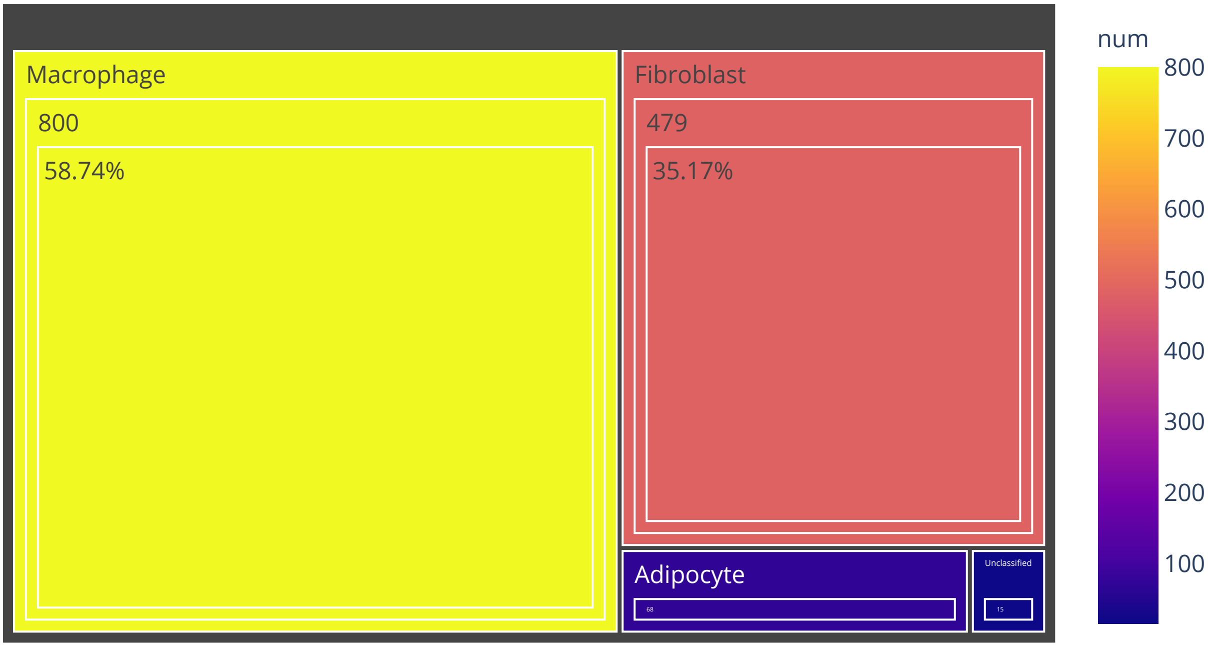

Organ:Pleura   Cell number:19695

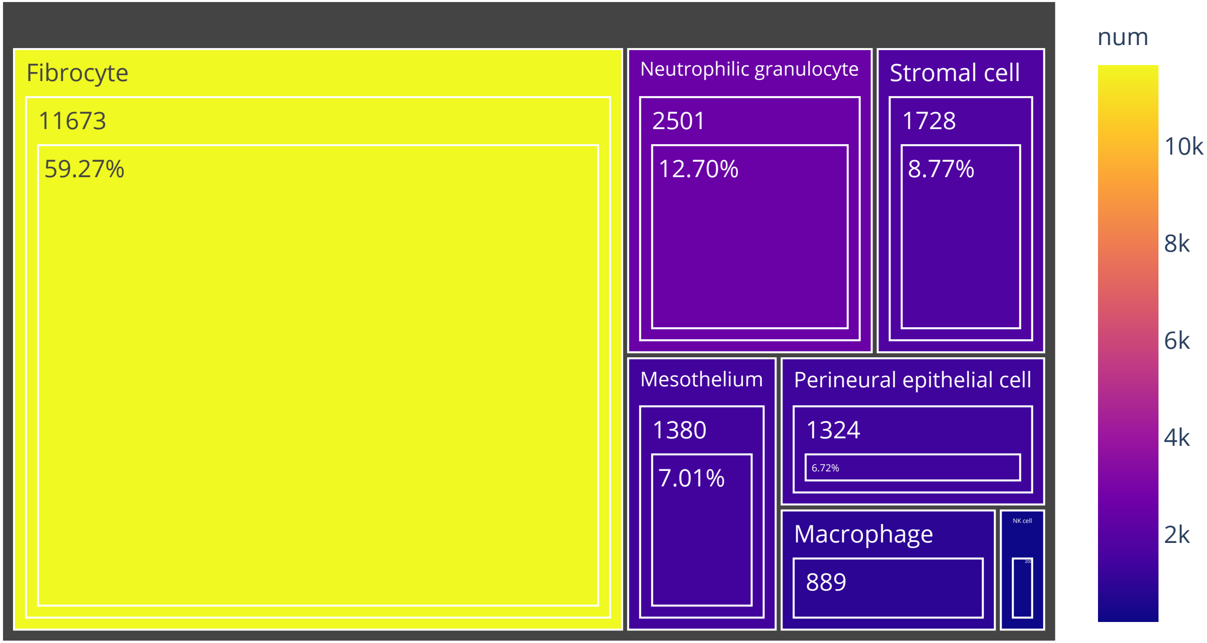

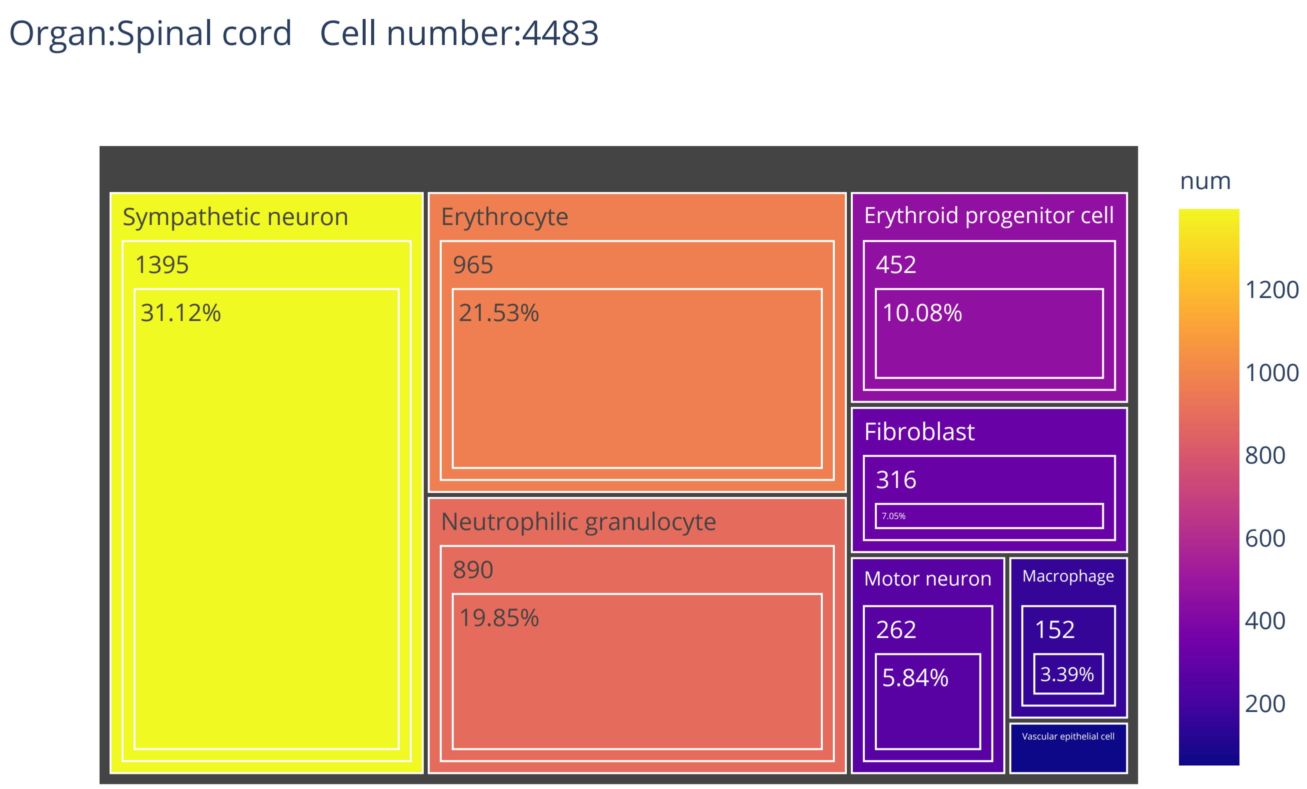

Organ:Spinal cord   Cell number:4483
Sympathetic neuron
1395
31.12%
Erythrocyte
965
21.53%
Neutrophilic granulocyte
890
19.85%
Erythroid progenitor cell
452
10.08%
Fibroblast
316
7.05%
Motor neuron
262
5.84%
Macrophage
152
3.39%
Vascular epithelial cell
num
1200
1000
800
600
400
200

# Organ:Intestine   Cell number:41851

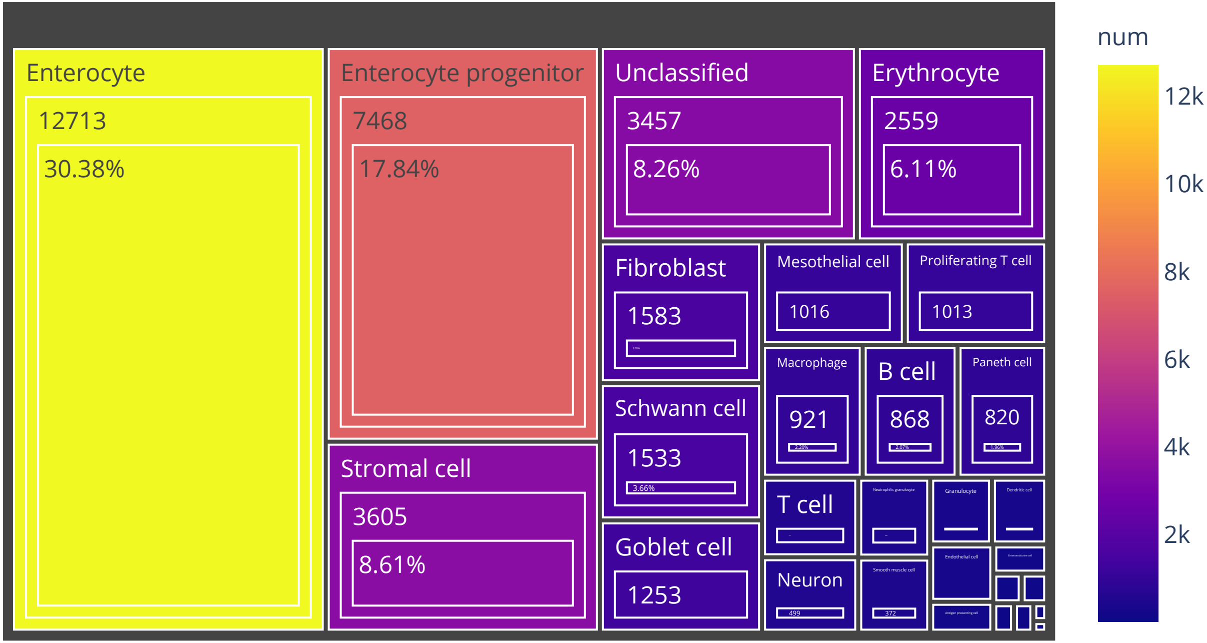

# Organ:Oesophagus   Cell number:87947

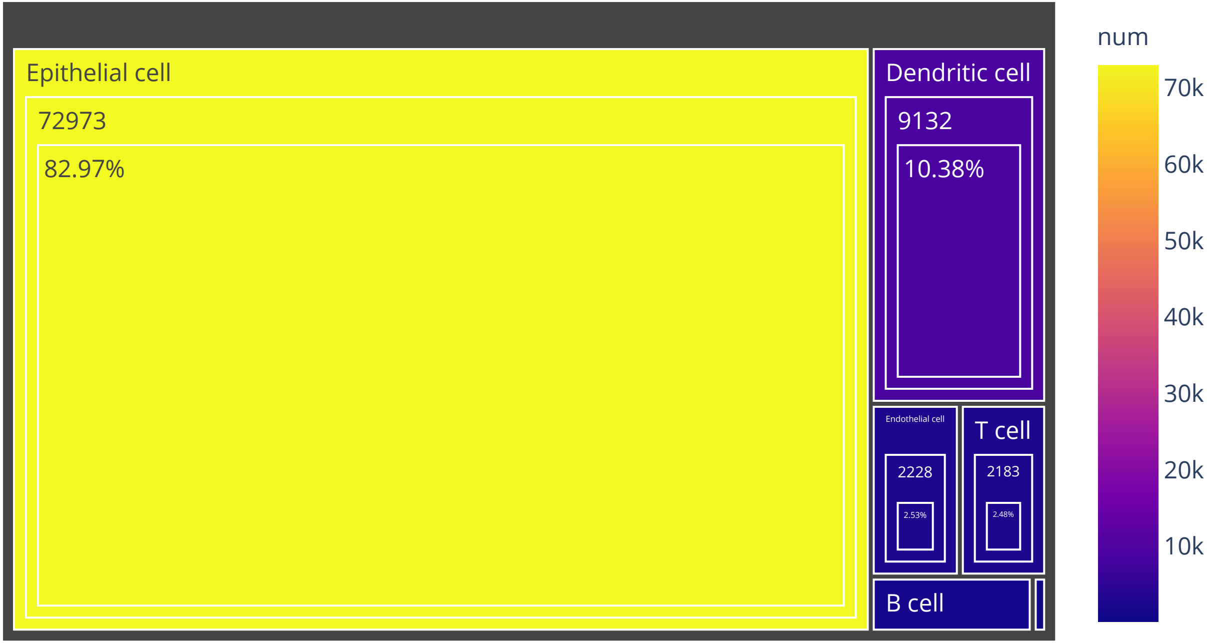

Organ:Bone marrow   Cell number:8671

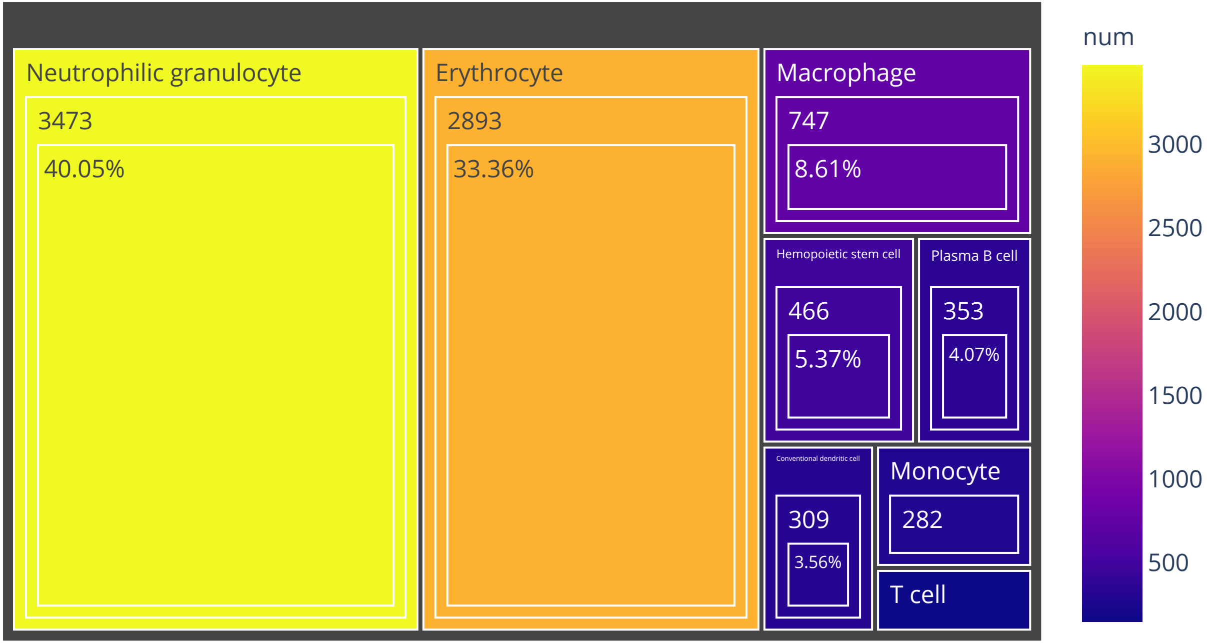

Organ:Thyroid   Cell number:12599

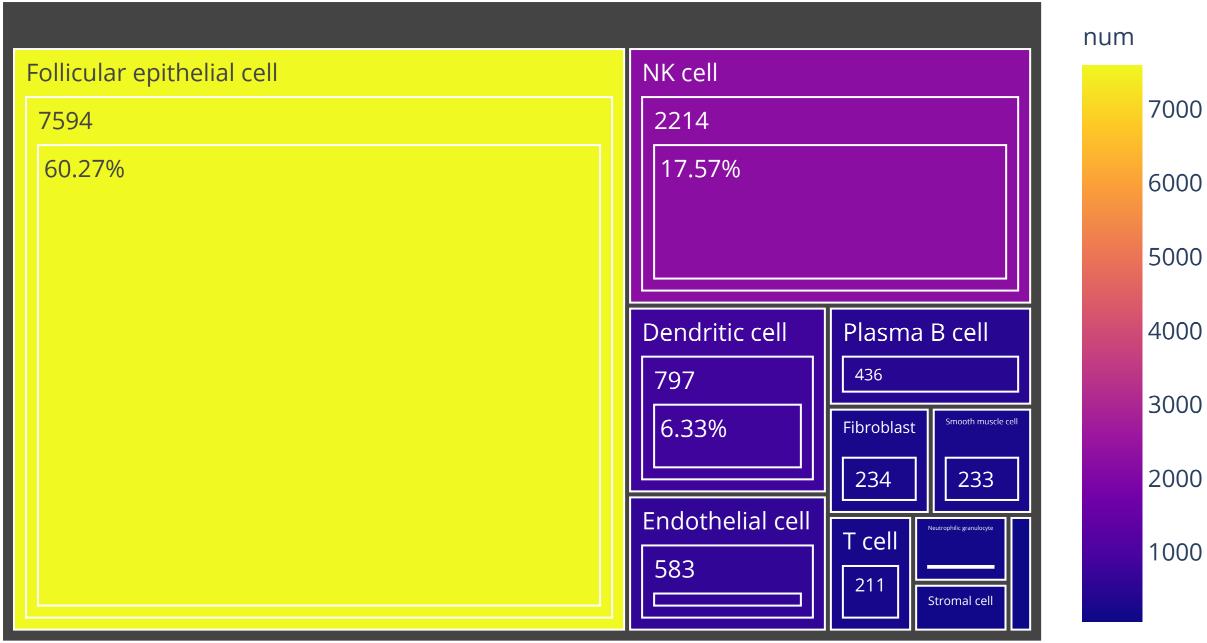

Organ:Vessel   Cell number:9652

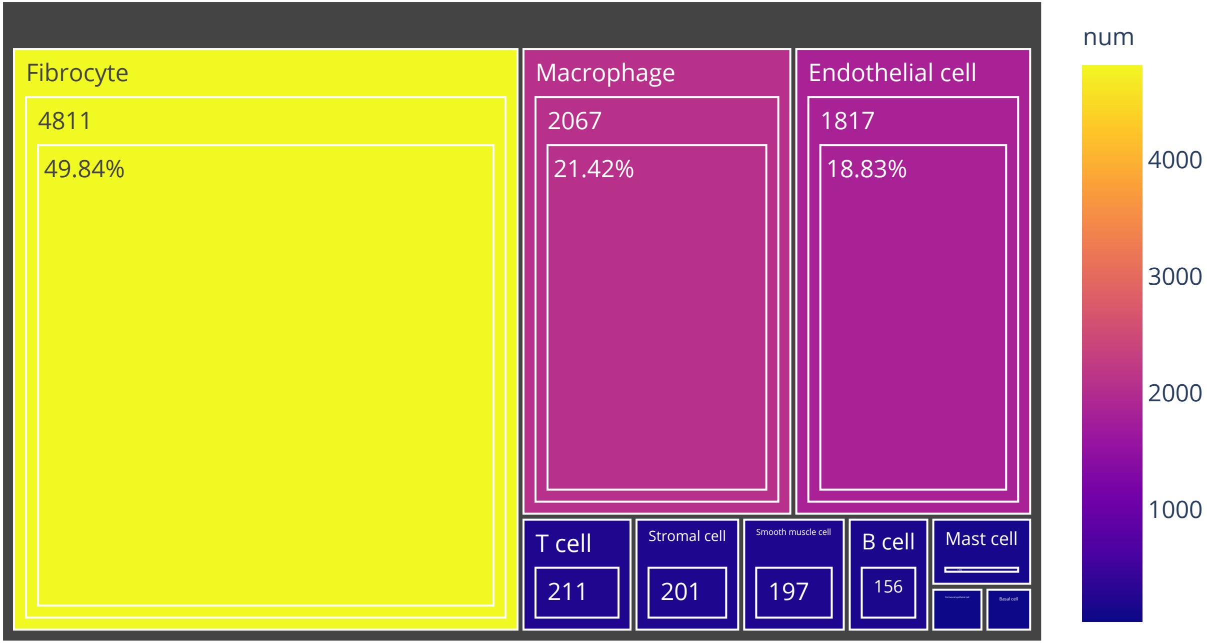

Organ:Kidney Cell number:43013

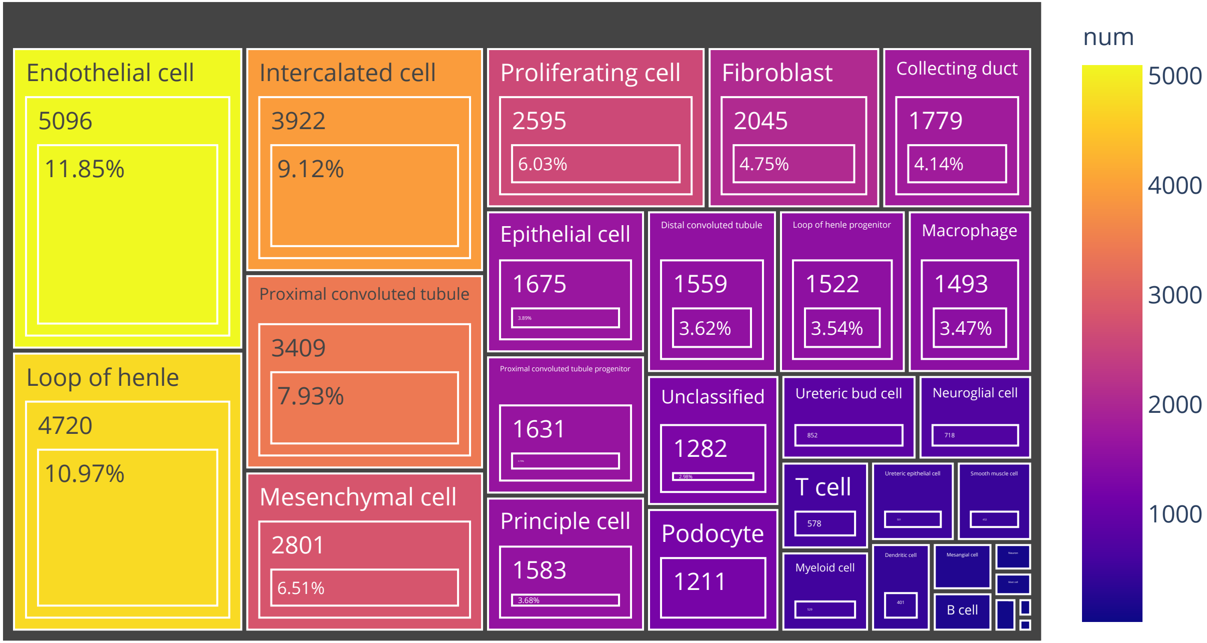

# Organ:Rib   Cell number:5907

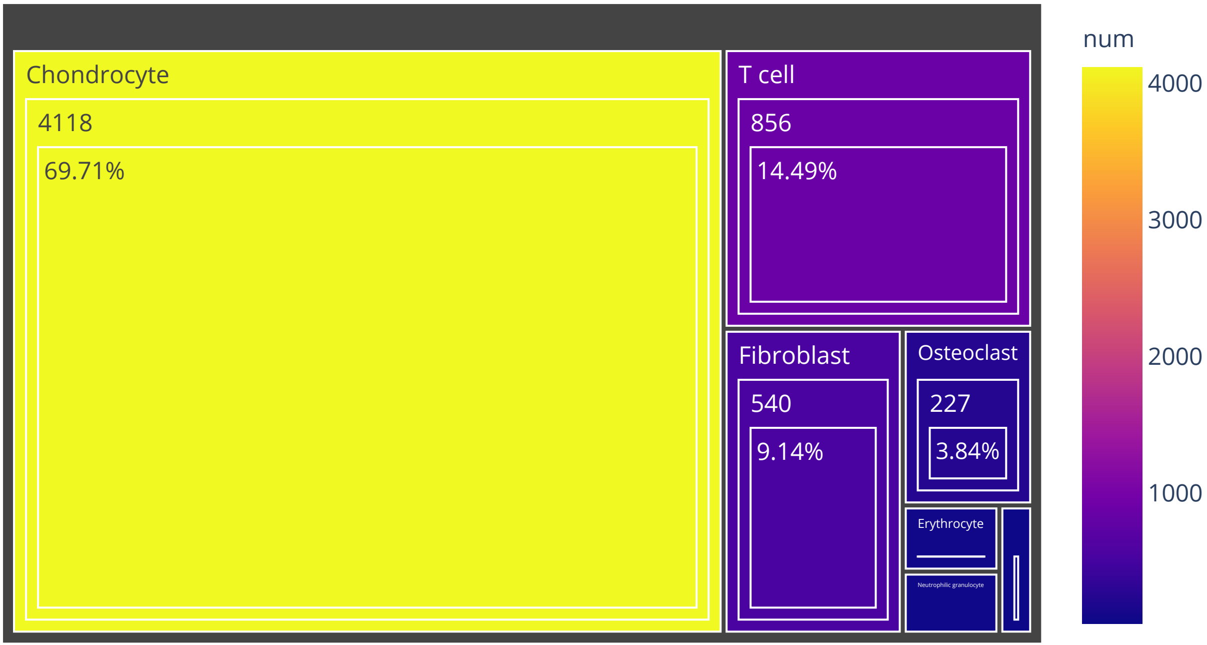

# Organ:Liver   Cell number:26475

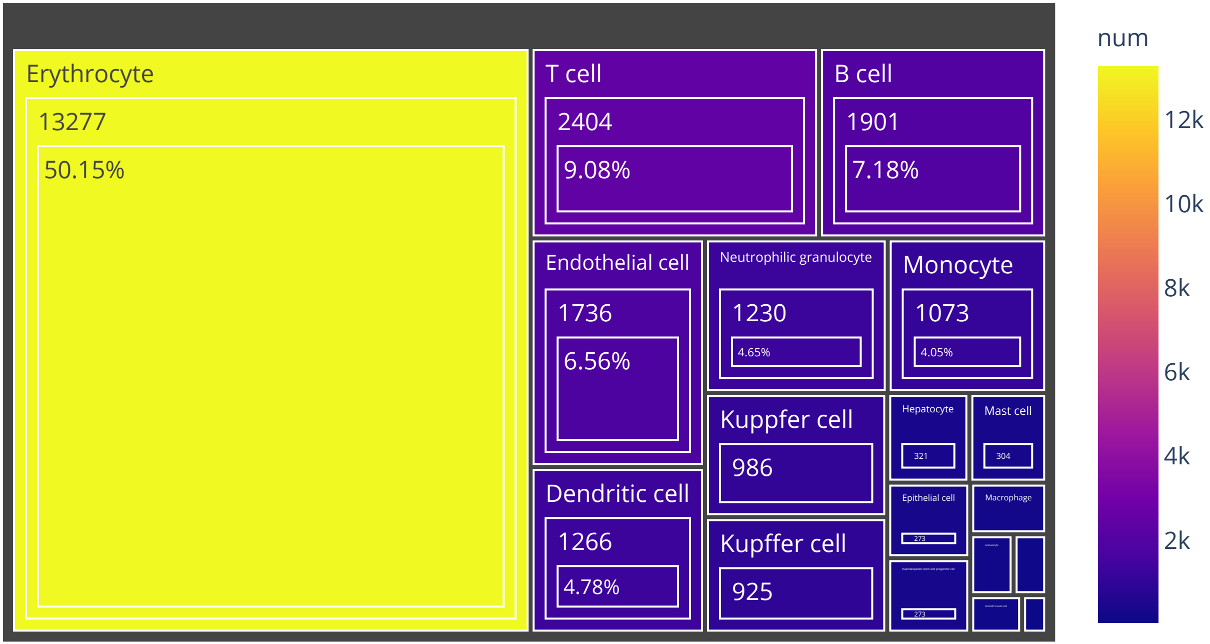

# Organ:Colon   Cell number:22919

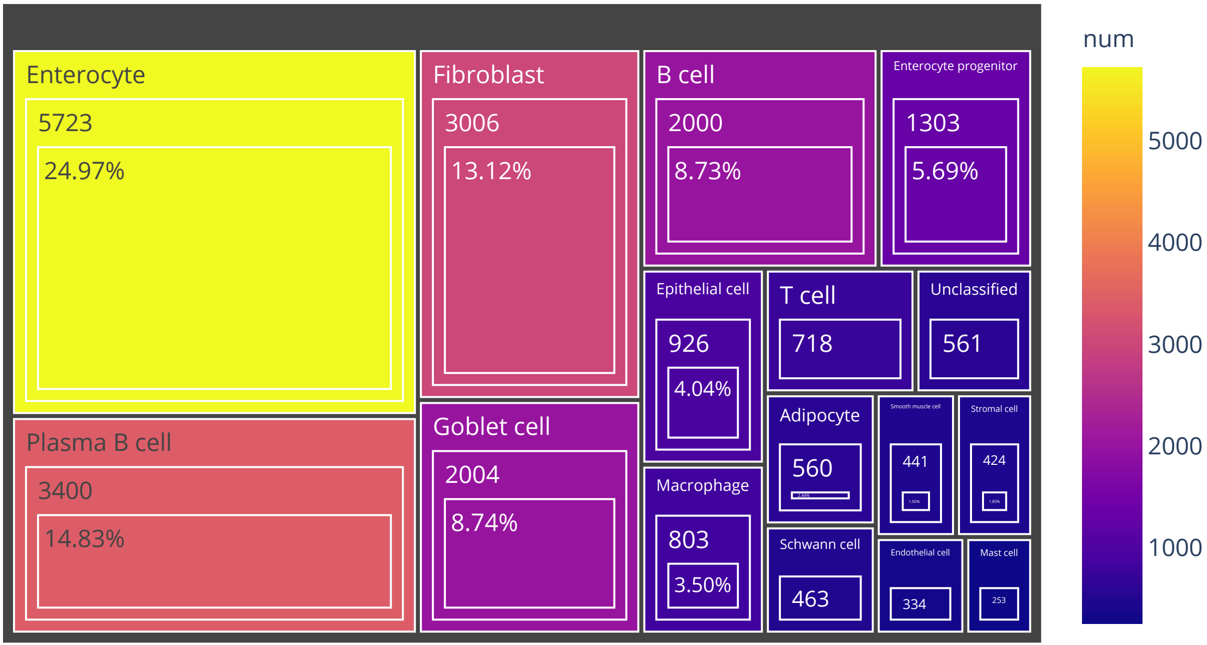

# Organ:Heart   Cell number:210597

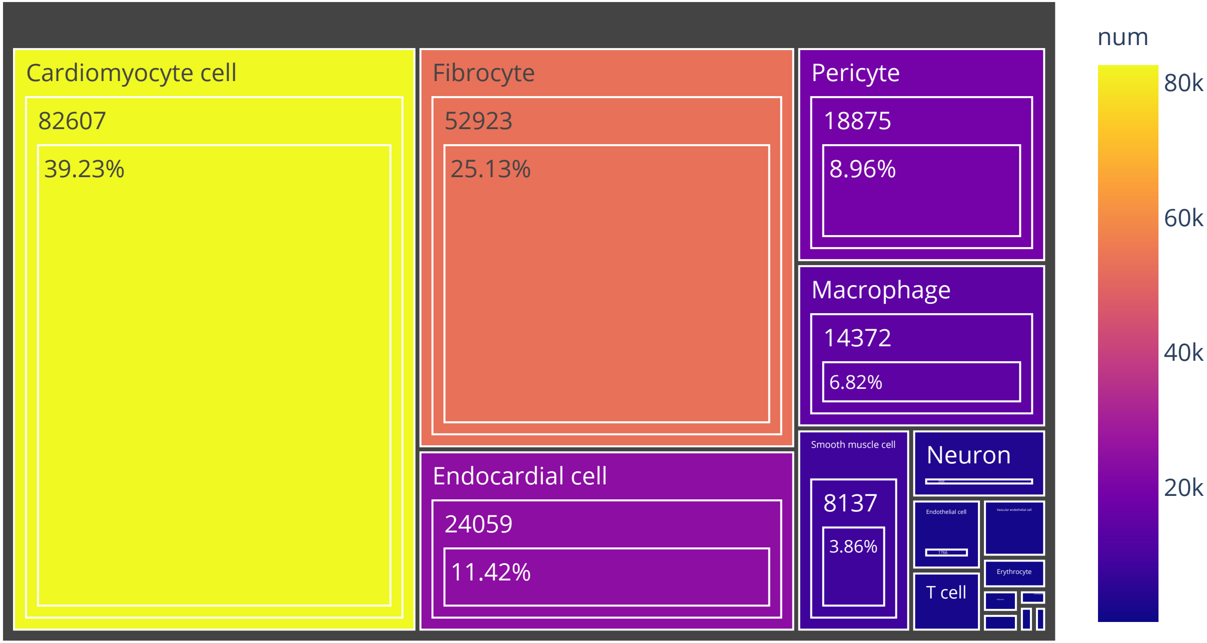

Organ:Gallbladder Cell number:14733

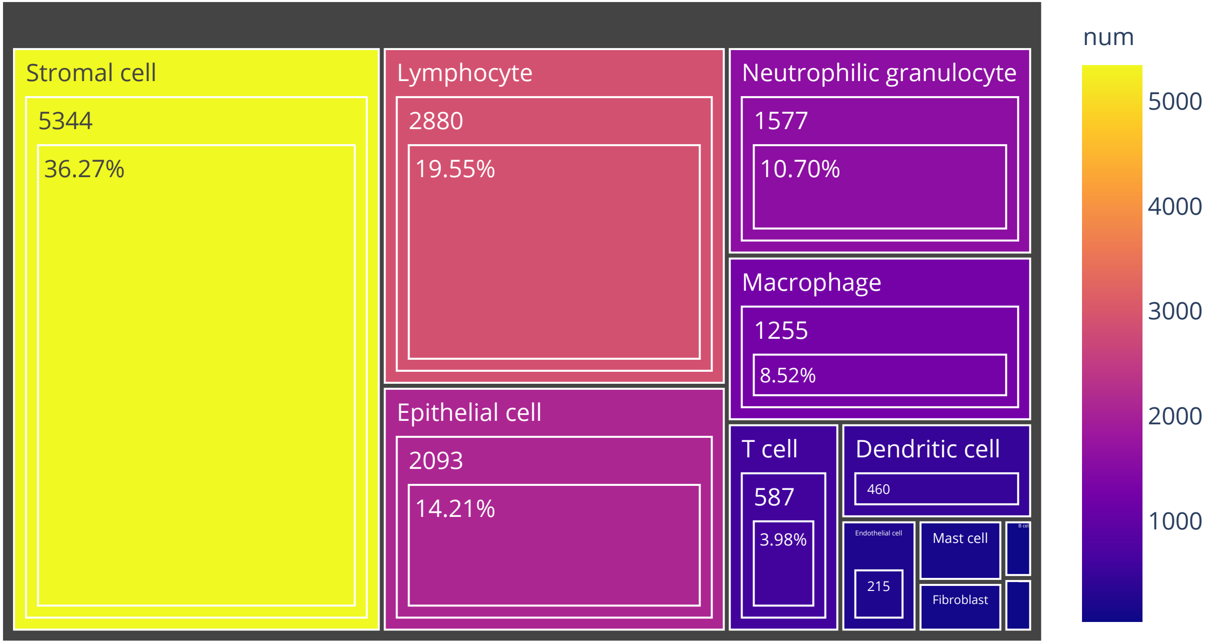

Organ:Muscle   Cell number:26029

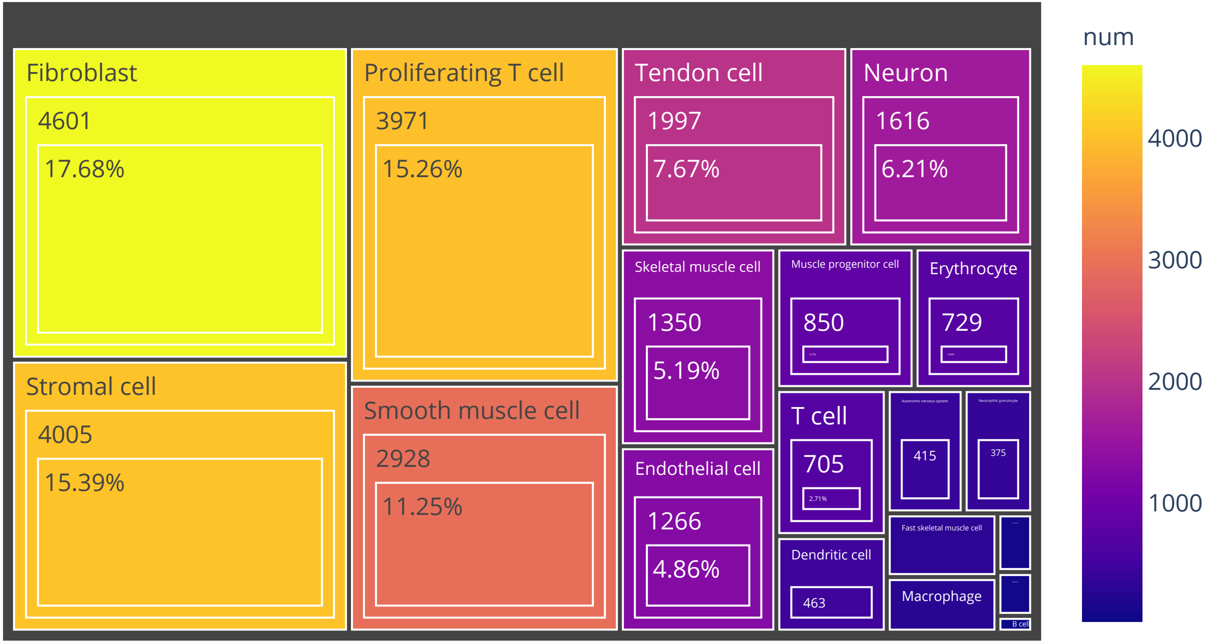

# Organ:Pancreas   Cell number:26566

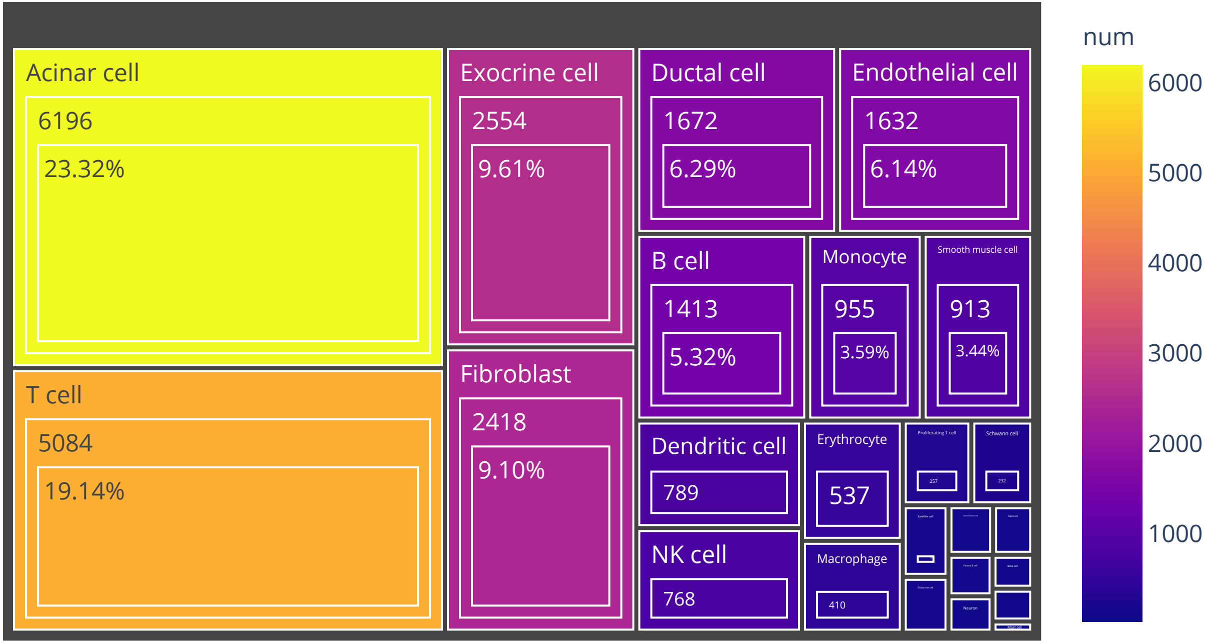

# Organ:Uterus   Cell number:8096

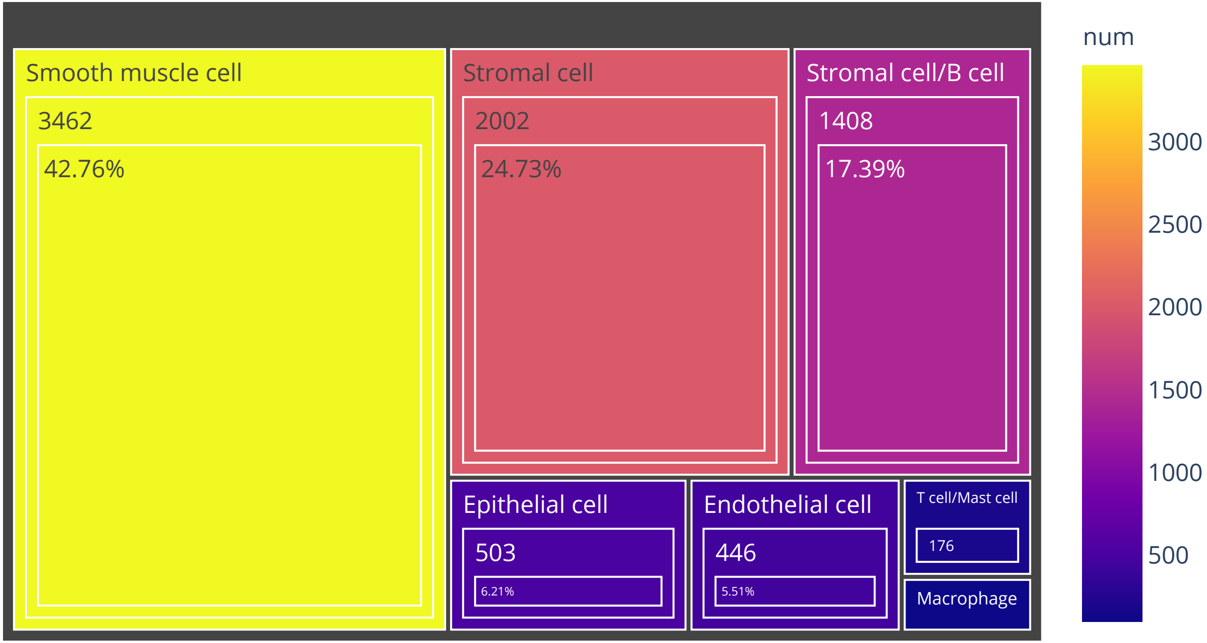
